# Tubulin autoregulation controls the biosynthesis of γ-tubulin to ensure mitotic fidelity

**DOI:** 10.64898/2026.01.09.698560

**Authors:** Maxim Assaf, Chahrazed Lacheheub, Ivana Gasic

## Abstract

Microtubule organization relies on the precise control of tubulin abundance to ensure accurate cytoskeletal function and faithful cell division. While α- and β-tubulin levels are controlled by a well-characterized autoregulatory pathway that triggers co-translational mRNA decay in response to excess soluble tubulin, how cells regulate the abundance of the core microtubule nucleator γ-tubulin has remained unclear. Here, we show that γ-tubulin is regulated by the canonical tubulin autoregulatory machinery. We find that γ-tubulin-encoding mRNAs are downregulated in response to elevated soluble αβ-tubulin levels. This regulation requires TTC5 to co-translationally recognize a conserved amino-terminal MPREI motif in nascent γ-tubulin proteins, and further recruit SCAPER and CCR4-NOT complex, targeting γ-tubulin mRNAs for decay. Disruption of this regulatory mechanism elevates γ-tubulin protein levels, increases centrosomal microtubule nucleation output, and compromises mitotic fidelity. Together, our findings establish γ-tubulin as a previously unrecognized substrate of tubulin autoregulation and reveal coordinated control of tubulin biosynthesis as a key mechanism for tuning microtubule nucleation and ensuring accurate chromosome segregation.

## Introduction

Microtubules are dynamic cytoskeletal filaments that maintain cell shape, position organelles, and direct intracellular transport [1]. During cell division, microtubules build the mitotic spindle that powers the division of the genetic material [2,3]. To accurately carry out these functions, cells must assemble and dynamically control the localization and mass of microtubules. The assembly and organisation of microtubules depend upon polymerization of the core building blocks—heterodimers of α- and β-tubulin proteins (henceforth αβ-tubulins) [4]. The rate-limiting step in microtubule formation is nucleation, mediated by γ-tubulin, a homolog of α- and β-tubulin [5–7]. Genomes of most vertebrates, including mammals, encode two γ-tubulin genes, TUBG1 and TUBG2 [8]. While TUBG1 is ubiquitously expressed, TUBG2 is neuron-specific and hardly detectable or absent in other cell types and tissues [9,10]. At microtubule-organizing centres (MTOCs) such as centrosomes [5,6], γ-tubulins together with gamma complex proteins (GCP2-6) form the γ-tubulin ring complex (γ-TuRC), a multi-subunit ring-shaped structure that templates microtubule nucleation [11–15].

Microtubule nucleation, polymerization, and depolymerization rates depend on the concentration of αβ-tubulins as their building blocks, and γ-tubulin as the primary nucleator, acting alongside additional context-specific nucleation factors [11,16–19]. Cells, therefore, must tightly control the production of tubulin proteins. For α- and β-tubulin, this is achieved via tubulin autoregulation, a co-translational mRNA degradation mechanism activated by excess soluble αβ-tubulins [20–23].

Through tubulin autoregulation, cells respond to excess unpolymerized αβ-tubulins by selectively degrading α- and β-tubulin–encoding mRNAs (TUBAs and TUBBs) [20–22]. The ribosome-associated factor TTC5 (Tetratricopeptide Repeat Protein 5) recognizes the nascent amino-terminal autoregulatory MREC and MREI motifs of α- and β-tubulins, respectively, as they emerge from the ribosome [22,23]. This amino-terminal motif confers specificity for TTC5 binding. Mutations at the second (arginine, R2) or third (glutamic acid, E3) residues of this motif abolish β-tubulin autoregulation [23]. Structural analyses rationalized these observations, showing that R2, E3, and I4 form key interactions with specific residues in the groove of TTC5 (R147, D225, and E259) [24]. Upon recognition of the nascent tubulin protein chain, TTC5 recruits the adaptor protein SCAPER (S-Phase Cyclin A–Associated Protein in the ER), which in turn engages the CCR4–NOT complex (Carbon Catabolite Repression–Negative on TATA-less) to promote mRNA decay via its CNOT11 (CCR4-NOT Transcription Complex Subunit 11) subunit [25].

By contrast, how cells regulate γ-tubulin abundance remains poorly understood, despite long-standing evidence that abnormal γ-tubulin levels disrupt the microtubule cytoskeleton and its functions [9,26–32]. Both overexpression [28,29] and depletion [30,31] of γ-tubulin have been shown to lead to dysfunctional mitotic spindles and impaired fidelity of mitotic chromosome segregation in various organisms. Mutations in monoubiquitination sites on γ-tubulin result in decreased γ-tubulin degradation and are associated with centrosome amplification and hyperactivity, highlighting the importance of quantity control [33,34].

Surprisingly, transcriptomic analyses reveal that, in parallel with TUBAs and TUBBs transcripts, cells downregulate the expression of γ-tubulin encoding mRNA (TUBGs) under microtubule-destabilizing conditions that elevate soluble αβ-tubulin levels, raising the possibility that γ-tubulin might also be subject to autoregulation [35]. These findings contrast with earlier studies that proposed γ-tubulin to employ a regulatory mechanism distinct from the one shared by α- and β-tubulins [29].

In this study, we investigated whether γ-tubulin is regulated by the canonical tubulin autoregulatory pathway, and how this regulation contributes to microtubule nucleation and mitotic fidelity in cultured human cells.

## Results

### Cells regulate TUBG expression in response to soluble αβ-tubulin levels

To determine whether γ-tubulin expression is regulated under conditions that destabilize or stabilize the microtubule network, we reanalyzed publicly available transcriptomic datasets from cancer cell lines treated with microtubule-targeting agents [36]. Differential expression analysis revealed that the ubiquitously expressed and abundant TUBG1 transcripts were significantly reduced in over 60% of cell lines exposed to eribulin, a microtubule-destabilizing agent (**Fig. 1A**). A similar trend was observed for TUBG2, although the changes in transcript levels were more variable across cell lines (**Fig. 1A**), likely due to its low abundance in non-neuronal cell types [9,37]. Conversely, treatment with paclitaxel, a microtubule-stabilizing drug, increased expression of both TUBG1 and TUBG2 in approximately half of the lines, with the remaining lines showing little or no response, suggesting that γ-tubulin transcript levels can depend on the microtubule polymerization state (**Fig. S1A**). This differential gene expression pattern in response to increased and decreased soluble αβ-tubulin levels is reminiscent of that of TUBAs and TUBBs, known to be post-translationally mediated via tubulin autoregulation [35].

**Figure 1.**
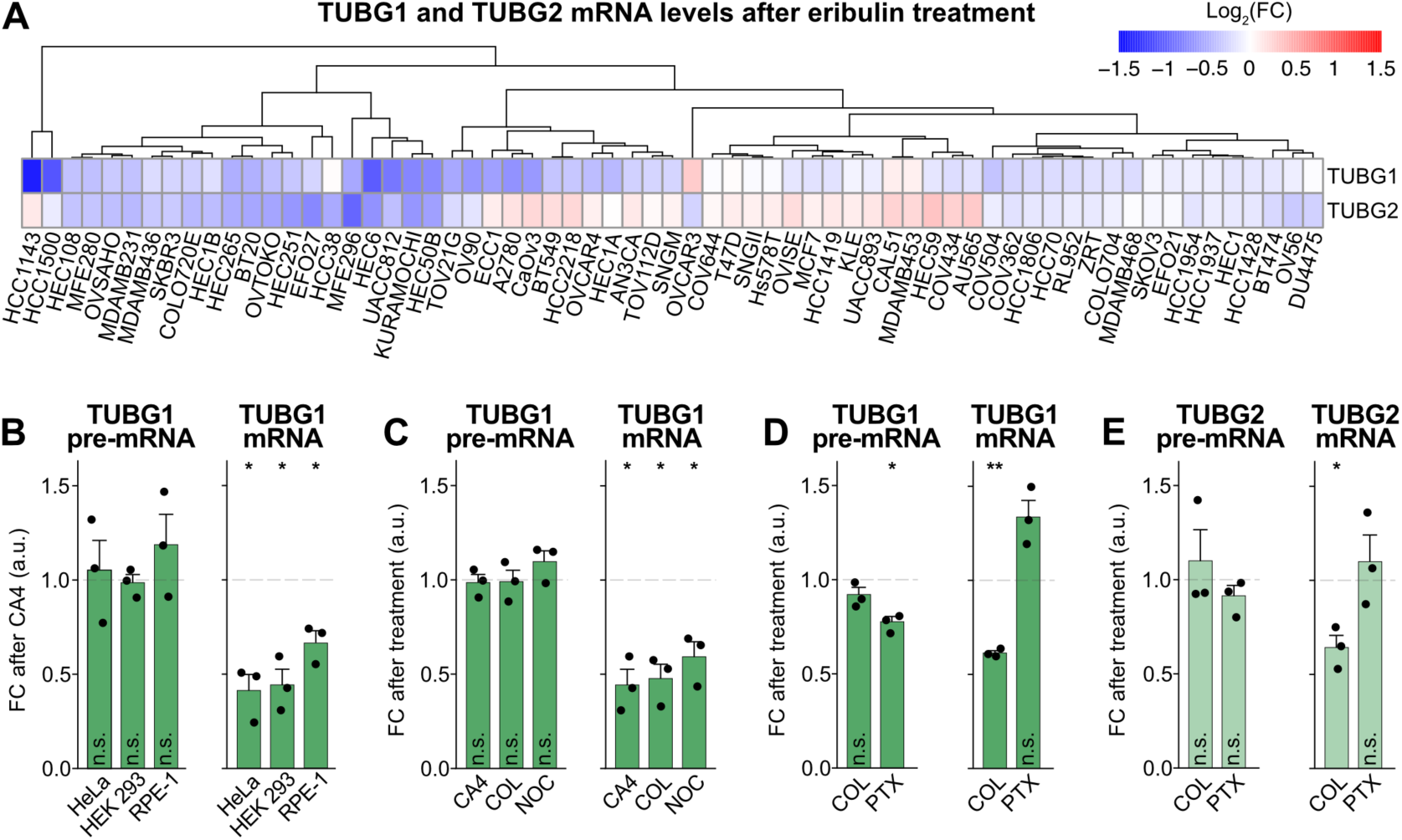
Cells regulate TUBG expression in response to soluble αβ-tubulin levels. (**A**) Relative expression levels of TUBG1 and TUBG2 in various cancer cell lines treated with eribulin for 24 h. Data were obtained from the GEO database (GEO series GSE50811, GSE50830, and GSE50831 [34]). Columns correspond to cells and rows to genes. Hierarchical clustering was performed on cells using Euclidean distance and complete linkage, and the results are displayed as a dendrogram above the heatmap, grouping cells with similar TUBG1/TUBG2 expression responses to treatment. Data are represented as Log_2_ Fold Change (Log_2_FC) relative to DMSO control, with the color key depicted on the right. (**B**) TUBG1 mRNA and pre-mRNA levels in HeLa, HEK293, and RPE-1 cell lines following 6 hours of treatment with CA4. (**C**) TUBG1 mRNA and pre-mRNA levels in HEK293 cells after 6 hours of treatment with the indicated small molecule inhibitors. (**D-E**) TUBG1 (D) and TUBG2 (E) mRNA and pre-mRNA levels in HeLa cells after 6 hours of treatment with either 100 nM COL or 300 nM PTX. All mRNA and pre-mRNA levels were normalized to the housekeeping transcripts RPL19 and GAPDH, respectively, and to DMSO-treated controls. Data represent mean ± SD from three independent experiments. A two-tailed Student’s *t*-test was performed for each of the cell lines with the DMSO-treated cells as reference. * *p* < 0.05, ** *p* < 0.01, *** *p* < 0.001, n.s. = not significant.

Reanalyzing publicly available transcriptomic data from rats exposed to various microtubule-destabilizing agents, we confirmed that this regulatory mechanism extends beyond cancer cell lines to living organisms (**Fig. S1B**) [38]. Consistent with the findings in eribulin-treated cancer cells, microtubule destabilization via vinca alkaloids or colchicine resulted in reduced TUBG1 mRNA levels in rat myocardium (**Fig. S1B**). Given that TUBA and TUBB mRNAs are known to undergo translation-dependent degradation in response to elevated soluble αβ-tubulin levels [24], we hypothesized that TUBG mRNA might be regulated through a similar pathway.

To investigate the mechanisms governing γ-tubulin expression, we simultaneously measured TUBG pre-mRNA and mRNA levels in control and cells treated with microtubule destabilizing poisons. This assay distinguishes transcriptional (pre-mRNA) from post-transcriptional (mRNA) regulation [33]. As previously reported for TUBB transcripts, in HeLa, HEK293, and RPE-1 cells, treatment with the microtubule destabiliser combretastatin A-4 (CA4) did not lead to any changes in unspliced TUBG1 pre-mRNA levels, indicating unaltered transcription (**Fig. 1B**). In contrast, spliced TUBG1 mRNA levels decreased following microtubule depolymerisation (**Fig. 1B**), mirroring differential expression of TUBB mRNAs (**Fig. S1C**) and suggesting post-transcriptional regulation of TUBG mRNAs. To rule out a drug-specific effect, we treated HEK293 cells with different microtubule-destabilizing agents, including colchicine (COL) and nocodazole (NOC). In line with previous results for TUBB transcripts (**Fig. S1D**), TUBG1 pre-mRNA remained intact, while the mRNA was downregulated upon the addition of all microtubule-destabilizing agents (**Fig. 1C**), confirming post-transcriptional regulation triggered by microtubule destabilization and not the inhibitor itself. Conversely, microtubule stabilization with paclitaxel (PTX) led to slightly decreased pre-mRNA, but increased TUBG1 mRNA levels (**Fig. 1D**), as seen for TUBBs (**Fig. S1E**). The regulation of TUBG2 mRNA mirrored that of TUBG1 (**Fig. 1E**). As TUBG2 is neuron-specific and hardly detectable in other cell types, we focused on TUBG1 for the remainder of this study [9].

Collectively, these data show that TUBG mRNA is regulated post-transcriptionally in response to changes in soluble αβ-tubulin levels, consistent with the tubulin autoregulatory mechanism.

### The TTC5-SCAPER-CCR4-NOT axis is required for γ-tubulin mRNA decay

Given their similar response to microtubule polymerization state, the stability of α-, β-, and γ-tubulin mRNAs may be controlled by the same tubulin autoregulatory machinery. To test this, we measured TUBG1 transcript levels in HeLa cells lacking key effectors of the tubulin autoregulation pathway (**Fig. 2A**). Following CA4 treatment, TUBG1 mRNA downregulation was abolished in TTC5 knockout (TTC5 KO) cells and restored by re-expression of wild-type TTC5^WT^ (**Fig. 2B**). Similarly, knockout of the adaptor protein SCAPER prevented TUBG1 mRNA reduction after CA4 treatment (**Fig. 2C**). TUBG1 mRNA downregulation was restored by the re-expression of SCAPER^WT^, but not by the autoregulation-defective SCAPER^Δ620^ mutant that fails to recruit the downstream effector CCR4-NOT complex (**Fig. 2C**) [25].

**Figure 2.**
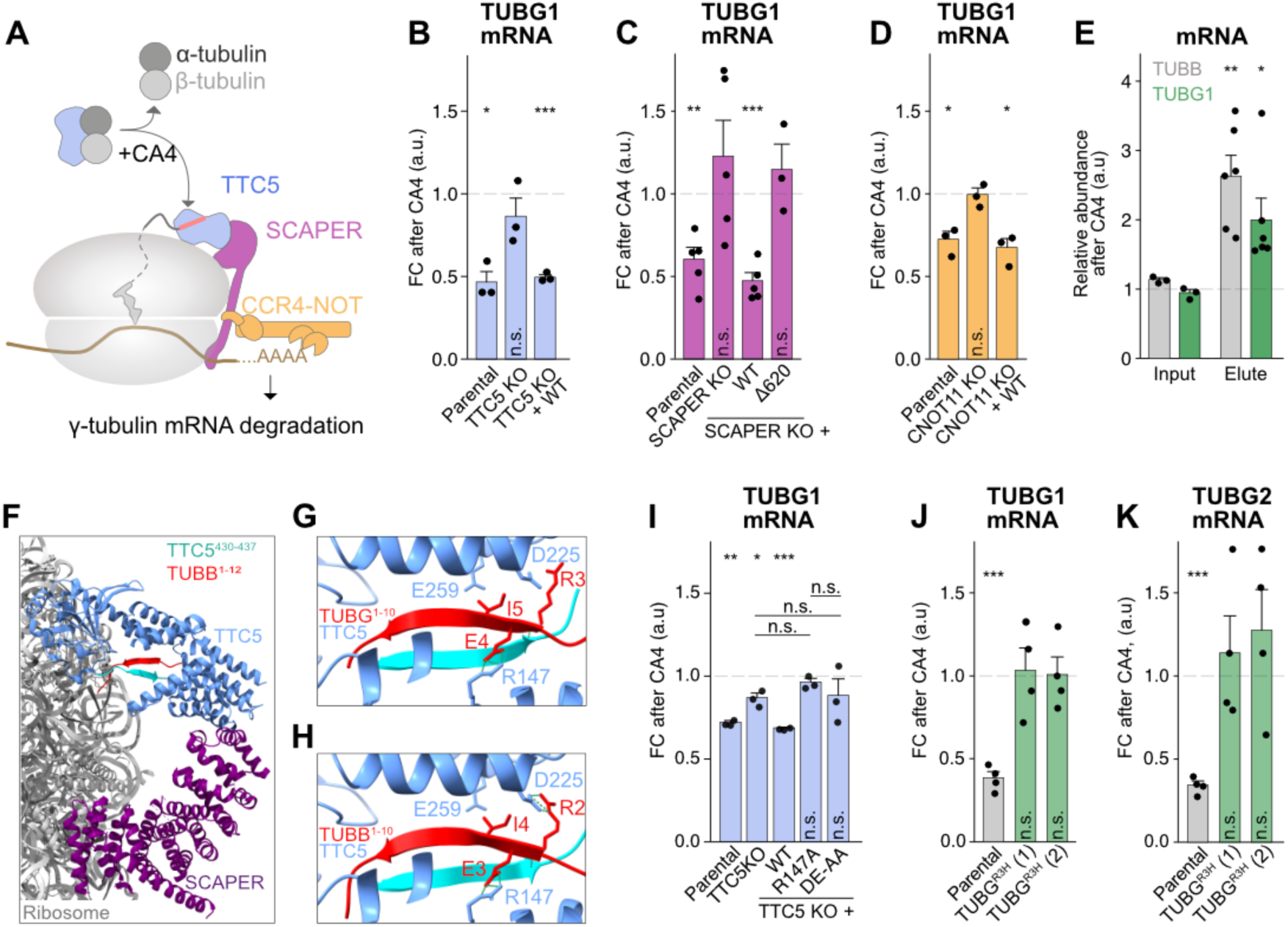
The TTC5-SCAPER-CCR4-NOT axis is required for γ-tubulin mRNA decay. (**A**) A hypothetical model for γ-tubulin mRNA regulation. (**B-D**) TUBG1 mRNA levels following 8-hour treatment with CA4 in HeLa cells of the indicated genotypes. Data represent mean ± SD with each data point corresponding to an independent experiment (n=3 for panels B and D, n=5 for panel C). Data were normalized to the housekeeping transcript RPL19. (**E**) β- and γ-tubulin mRNA levels before (input) and after (elute) co-immunoprecipitation of HEK293 lysates with recombinant Strep-TTC5. Data are normalized to housekeeping transcript RPLP1 (mean ± SD; n=3 input, n=6 elute). (**F**) AlphaFold2-predicted structure of the carboxy-terminal domain of TTC5 (cyan) interacting with the nascent β-tubulin (red), fitted into the published cryo-EM density (PDB: 8BPO). (**G-H**) Close-up views of the AlphaFold3-predicted interactions between the carboxy-terminal domain of TTC5 and γ-tubulin (G) or β-tubulin (H) nascent chains. Key residues in TTC5 and nascent tubulins are highlighted. (**I**) TUBG1 mRNA levels in TTC5 knockout and the indicated rescue cell lines following 8-hour treatment with 1 μM CA4. Data represent mean ± SD of three independent replicates. (**J-K**) TUBG1 (J) and TUBG2 (K) mRNA levels in TUBG^R3H^ cells following CA4 treatment. Data represent mean ± SD from four independent replicates, normalized to housekeeping transcript RPLP1. A two-tailed Student’s *t*-test was performed for each of the cell lines with the DMSO-treated cells as reference. * *p* < 0.05, ** *p* < 0.01, *** *p* < 0.001, n.s. = not significant.

Previous studies established CNOT11 as the specificity subunit required for CCR4-NOT-mediated TUBA and TUBB mRNA deadenylation and decay in tubulin autoregulation [25]. Consistent with the hypothesis that TUBG mRNAs are subject to tubulin autoregulation, CNOT11 KO cells were also unable to decrease TUBG1 mRNA levels following microtubule destabilization (**Fig. 2D**). This was reversed by the re-expression of CNOT11^WT^ (**Fig. 2D**), confirming CCR4-NOT-mediated downregulation. As expected, no changes in transcriptional regulation of TUBG1 were observed in any of the knockout cell lines, consistent with the post-transcriptional and co-translational nature of the autoregulation pathway (**Fig. S2A–C**). Together, these data reveal TUBG mRNA regulation via the TTC5-SCAPER-CCR4-NOT axis, suggesting that nascent γ-tubulins are recognized by TTC5 as they emerge from the ribosome.

### TTC5 cotranslationally recognizes the γ-tubulin MPREI amino-terminal motif

The amino-terminal peptides of nascent α- and β-tubulins confer specificity for TTC5 binding. However, the γ-tubulin amino-terminal sequence (MPREI) differs from that of α- and β-tubulins (MREC/MREI) by the insertion of a proline residue in the second position, shifting the critical arginine and glutamic acid residues to positions three and four, respectively (**Fig. S3A**). To assess whether the proline insertion is permissive for TTC5 binding on γ-tubulin mRNA-translating ribosomes, we performed selective pulldown of recombinant TTC5 added to lysates from TTC5 KO cells pre-treated with DMSO (control) or CA4. As expected, in DMSO-treated lysates, recombinant TTC5 was sequestered by αβ-tubulin, preventing its translocation to ribosomes (**Fig. S3B**) [22]. This sequestration was alleviated by CA4 treatment before cell lysis, allowing TTC5 to engage ribosomes via emerging nascent tubulin chains, as evidenced by the presence of ribosomal proteins in the pulldown (**Fig. S3C**). Quantitative analysis of RNA eluted from TTC5 pulldowns showed significant enrichment of TUBG1 and TUBB mRNAs following CA4 treatment (**Fig. 2E**), indicating that TTC5 can recognise and bind to γ-tubulin nascent chains emerging from the ribosomes.

These data suggest that, despite the proline insertion in the second position, TTC5 can accommodate the MPREI motif of γ-tubulin within its binding groove. Consistent with this, an AlphaFold3 structural prediction shows that TTC5 houses the key arginine and glutamic acid residues (R3 and E4) of γ-tubulin in the same configuration as R2 and E3 of β-tubulin (**Fig.2F-H and S3D-F**) [39]. Consequently, the γ-tubulin nascent chain appears to be positioned deeper into the TTC5 pocket (**Fig. 2F-G**). Notably, the key electrostatic interactions between the R147, D225, and E259 of TTC5 and R3, E4, and I5 of MPREI nascent peptide are predicted to be maintained (**Fig. 2G**). Supporting this model, cells expressing TTC5 where R147, or D225 and E259 residues are mutated to alanine (TTC5^R147A^ and TTC5^DE-AA^, respectively) lose the ability to autoregulate TUBG mRNA upon microtubule destabilization (**Fig. 2I**), as seen for TUBB mRNAs (**Fig. S3G**).

Based on these observations, we hypothesized that mutating the MPREI motif of γ-tubulin would disrupt the regulation of its mRNA. However, as the amino-terminus of γ-tubulin lies at the interface with GCPs [40–42], extensive alterations could compromise its structural integrity or function. To minimize this risk while testing the regulatory mechanism, we minimally mutated the γ-tubulin amino-terminal sequence, substituting arginine 3 with histidine—an alteration previously shown to be sufficient to abolish tubulin autoregulation in β-tubulin [23, 24].

Using CRISPR-Cas9-mediated genome editing, we generated HeLa cell lines expressing γ-tubulin^R3H^ (TUBG1^R3H^) from the endogenous *TUBG1* locus (**Fig. S4A**). In these cells, TUBG1 mRNA decay was abolished upon microtubule destabilization (**Fig. S4B**). Next, we edited the endogenous *TUBG2* locus in the TUBG1^R3H^ background, generating HeLa cell lines expressing γ-tubulin^R3H^ from both *TUBG1* and *TUBG2* loci (TUBG^R3H^) (**Fig. S4C-D**). As expected, downregulation of both TUBG1 and TUBG2 mRNA in response to microtubule destabilization was abolished in TUBG^R3H^ cell lines (**Fig. 2J-K**). This effect was specific to TUBG mRNA, as TUBB mRNAs were efficiently degraded in response to microtubule destabilisation in these cells (**Fig. S4E**). Importantly, no significant changes were observed in TUBGCP2 and TUBGCP4 mRNA levels in wild-type parental cells, TTC5 KO, and TUBG^R3H^ cells treated with CA4, indicating that tubulin autoregulation acts specifically on γ-tubulin rather than the γ-TuRC complex (**Fig. S4F-G**).

Together, these results demonstrate that the γ-tubulin MPREI amino-terminal motif is required for TUBG mRNA regulation, consistent with TTC5 recognition. The R3H substitution disrupts this regulatory pathway, thereby providing a genetically stable system to dissect the functional significance of γ-tubulin mRNA regulation.

### Loss of TUBG mRNA regulation elevates γ-tubulin protein levels and centrosomal microtubule nucleation

To study the implications of deregulated γ-tubulin biosynthesis, we first probed the localisation and interactome of γ-tubulin^R3H^. Using an overexpression system, we verified that γ-tubulin^R3H^ retained centrosomal localisation in HeLa cells (**Fig. 3A**). Moreover, γ-tubulin^R3H^ co-immunoprecipitated with GCPs (**Fig. 3B**), indicating that the R3H mutation does not perturb γ-tubulin’s canonical localisation or interaction with the γ-TuRC scaffold. Finally, to distinguish between phenotypes that arise due to loss of γ-tubulin mRNA regulation and potential effects of the R3H mutation itself, independent of γ-tubulin mRNA regulation, we generated a γ-tubulin^R3H^ mutant in the TTC5 knockout background (TTC5 KO + TUBG^R3H^) (**Fig. S5A**).

**Figure 3.**
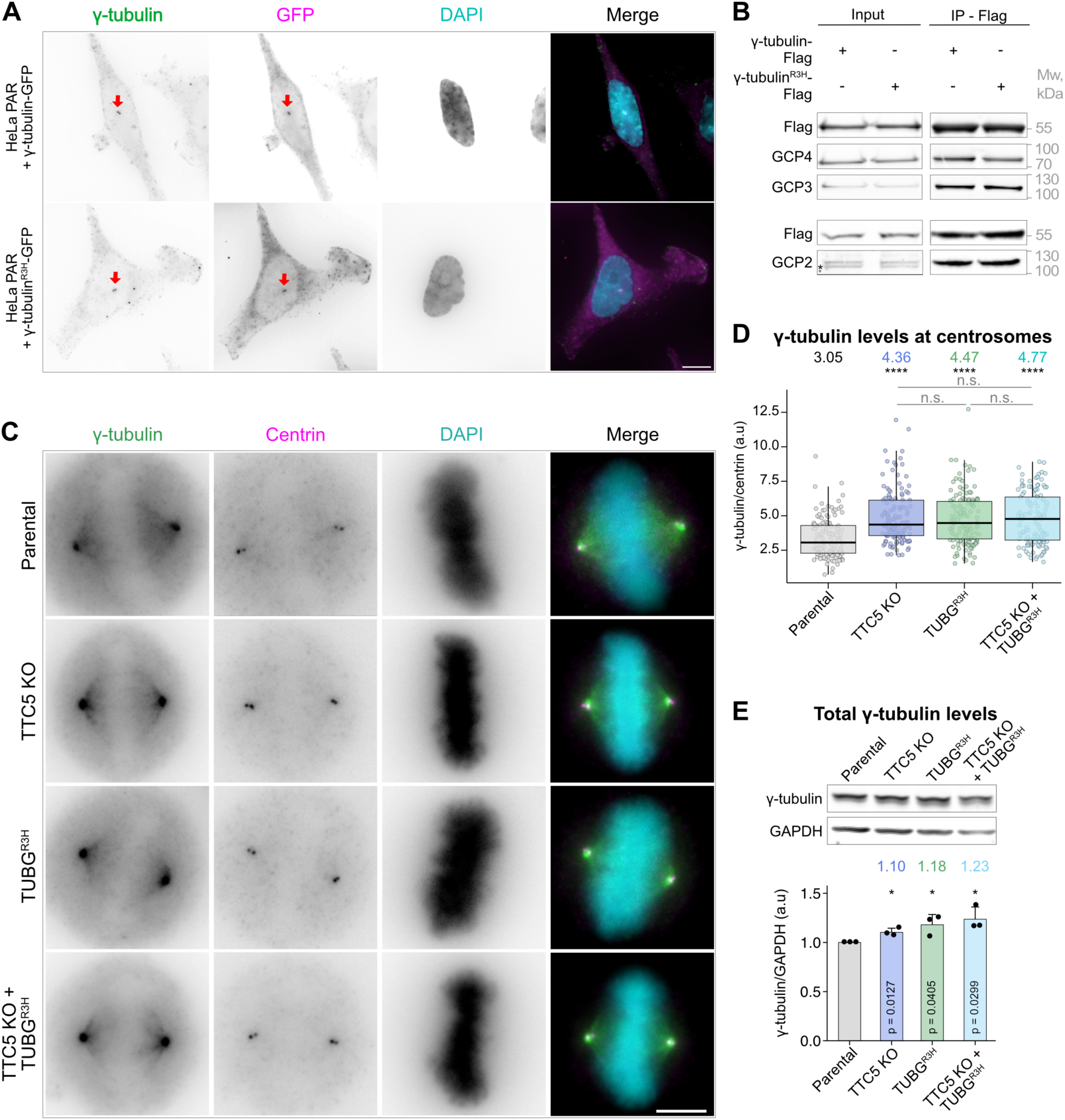
Loss of TUBG mRNA regulation elevates γ-tubulin protein levels. (**A**) Maximum intensity projections of HeLa parental cells stably overexpressing wild-type γ-tubulin-eGFP or γ-tubulin^R3H^-eGFP, immunostained for γ-tubulin (green) and GFP (magenta). Scale bar = 10 µm. (**B**) Indicated γ-tubulin-Flag constructs were stably expressed in HeLa parental cells using the FLP-FRT system and pulled down via the Flag-tag. Bound γ-tubulin complex components GCP2, GCP3, and GCP4 were visualized by western blot. * Annotates an unspecific band. (**C**) Representative images of mitotic cells immunostained for γ-tubulin (green) and centrin (magenta). (**D**) Quantification of centrosomal γ-tubulin fluorescence intensity normalized to centrin across the indicated cell lines. The experiment was conducted with three biological replicates, analysing 131, 129, 136, and 121 centrosomes for parental, TTC5 KO, TUBG^R3H^, and TTC5 KO + TUBG^R3H^ cells, respectively. Scale bar = 5 µm. Data show median ± interquartile range. Median values colour-coded by genotype are indicated above the respective boxplots. Shown are Benjamini-Hochberg-adjusted *p-*values from an unpaired Mann-Whitney test comparing ranks for each of the indicated cell lines with the parental cell line as reference. (**E**) Western blot analysis of total γ-tubulin protein levels in the indicated cell lines. Quantification shows γ-tubulin intensity normalized to GAPDH loading control and parental cell line. Data display mean ± SD from three independent experiments. Shown are *p-*values from an unpaired, two-tailed Student’s t-test comparing each cell line to the parental. * *p* < 0.05, ** *p* < 0.01, *** *p* < 0.001, **** *p* < 0.0001, n.s. = not significant.

Having established that γ-tubulin^R3H^ disrupts TTC5-mediated mRNA regulation while maintaining normal localization and γ-TuRC interactions, we examined how loss of this regulatory control affects γ-tubulin levels at the centrosome during cell division. Immunofluorescence staining of metaphase cells revealed elevated centrosomal γ-tubulin in TTC5 KO and TUBG^R3H^ cells (**Fig. 3C-D and S3B**). The increase in centrosomal γ-tubulin levels was phenocopied in TTC5 KO + TUBG^R3H^ cells (**Fig. 3C-D**), while centrin levels remained unaffected (**Fig. S5C**). We speculated that this increase in centrosomal γ-tubulin might reflect a global rise in γ-tubulin protein levels. Western blot analysis confirmed a moderate but consistent increase in γ-tubulin levels in TTC5 KO, TUBG^R3H^, and TTC5 KO + TUBG^R3H^ cells relative to wild-type parental cells (**Fig. 3E)**.

During interphase, the majority of γ-tubulin resides in the cytoplasm, with a smaller fraction localising to the centrosome [43–45]. At the onset of mitosis, γ-tubulin is rapidly and extensively recruited to the pericentriolar material (PCM) to mediate microtubule nucleation at the centrosome [43]. We reasoned that elevated γ-tubulin levels at the centrosomes of mutant cells may enhance their microtubule nucleating capacity. To test this, we assessed microtubule nucleation in mutant cells using a microtubule regrowth assay in metaphase cells. In this assay, microtubules are first depolymerised by cold treatment and then regrown upon rewarming, enabling visualisation of early stages of microtubule renucleation (**Fig. 4A and S6A**). We quantified centrosomal polymer mass within 45s of rewarming, a window enriched for nascent microtubule formation at centrosomes (**Fig. S6B**). As expected, TTC5 KO, TUBG^R3H^, and TTC5 KO + TUBG^R3H^ cells all showed a significant increase in α-tubulin fluorescence intensity at the centrosome shortly after regrowth, consistent with increased nucleation output (**Fig. 4B-C and S6C**).

**Figure 4.**
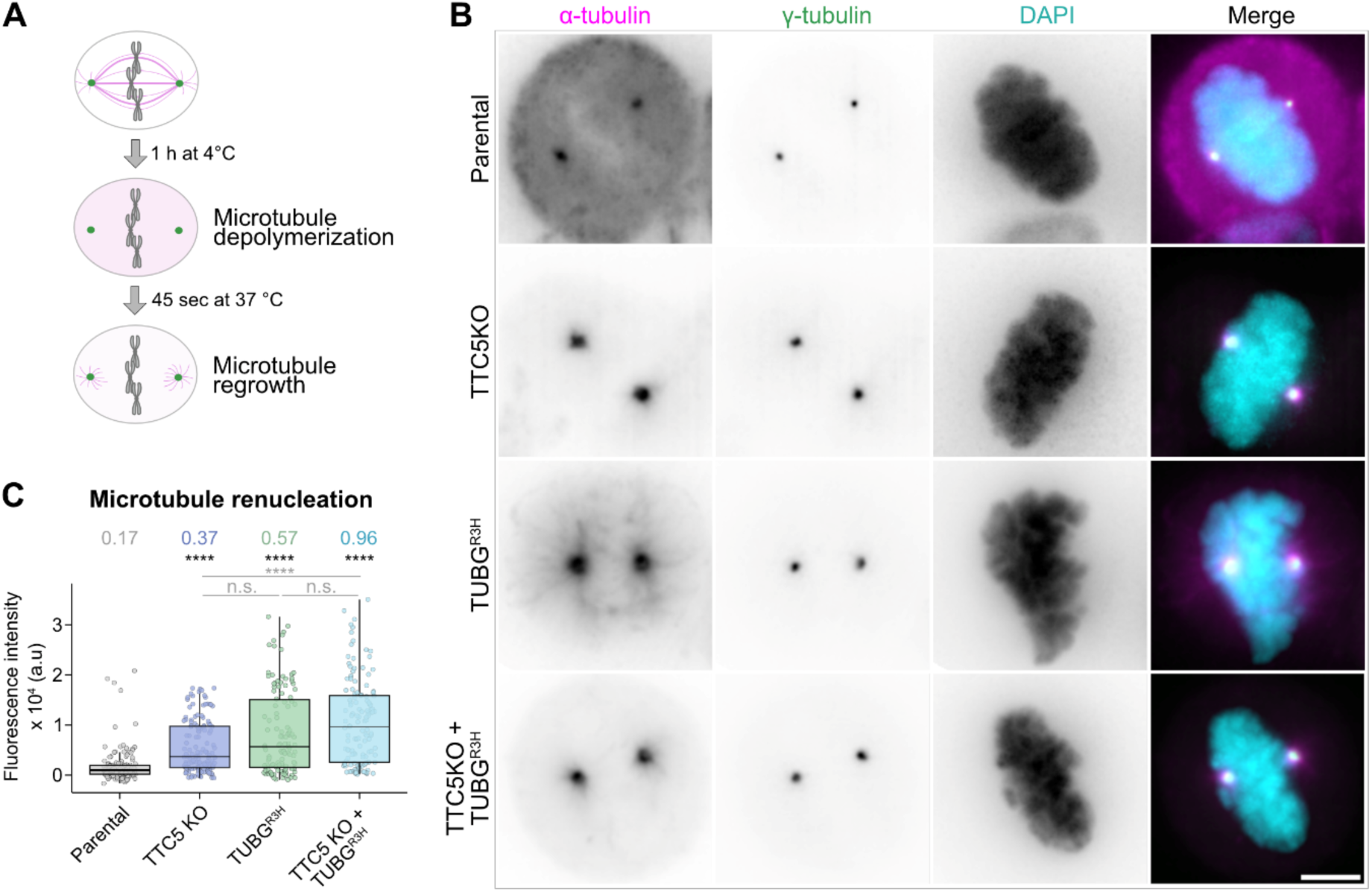
Loss of TUBG mRNA regulation elevates centrosomal microtubule nucleation output. (**A**) Schematic representation of the microtubule regrowth assay. (**B**) Maximum intensity projections of representative images of mitotic HeLa parental, TTC5 KO, TUBG^R3H^, and TTC5 KO + TUBG^R3H^ cells following cold-induced microtubule depolymerisation and 45 seconds of microtubule regrowth. Cells were immunostained for γ-tubulin (green) and α-tubulin (magenta). Scale bar = 5 µm. (**C**) Quantification of microtubule renucleation represented as α-tubulin fluorescence intensity. The experiment was done with four biological replicates, analysing 143, 118, 137, and 134 centrosomes for parental, TTC5 knockout, TUBG^R3H^, and TTC5 knockout + TUBG^R3H^ cells, respectively. Data show the median ± interquartile range. Median values colour-coded by genotype are indicated above the respective boxplots. Shown are Benjamini-Hochberg-adjusted *p-*values from an unpaired Mann-Whitney test comparing ranks for each of the indicated cell lines with the parental cell line as reference. **** *p* < 0.0001, n.s. = not significant.

Collectively, these results indicate that loss of γ-tubulin mRNA regulation moderately increases global γ-tubulin levels and centrosomal γ-tubulin accumulation during mitosis, increasing microtubule nucleation capacity.

### Regulated γ-tubulin biosynthesis maintains mitotic fidelity

Disruption of tubulin autoregulation has been shown to impair faithful chromosome segregation during mitosis [22,24,25]. Likewise, perturbations of γ-tubulin quantity led to mitotic errors [9,28–32]. We therefore asked whether γ-tubulin mRNA regulation contributes to mitotic accuracy and monitored chromosome segregation using live-cell imaging (**Fig. 5A**). In agreement with previous reports [22,24,25], TTC5 KO cells displayed a higher incidence of errors in chromosome alignment on the metaphase plate (2.4-fold) and segregation in anaphase (2.2-fold) compared to wild type parental cells (**Fig. 5B-D**). TUBG^R3H^ and TTC5 KO + TUBG^R3H^ cells also exhibited increased chromosome alignment errors (1.4- and 1.6-fold, respectively) relative to wild type parental cells (**Fig. 5B and 5C**). Notably, both TUBG^R3H^ and TTC5 KO + TUBG^R3H^ lines phenocopied the segregation defects observed in TTC5 KO cells, showing a 1.9-fold and 2.2-fold increase in anaphase errors, respectively (**Fig. 5B and 5D**). Despite decreased mitotic fidelity, γ-tubulin mRNA regulation-deficient cells progressed through mitosis with unaltered timing (**Fig. S7A**).

**Figure 5.**
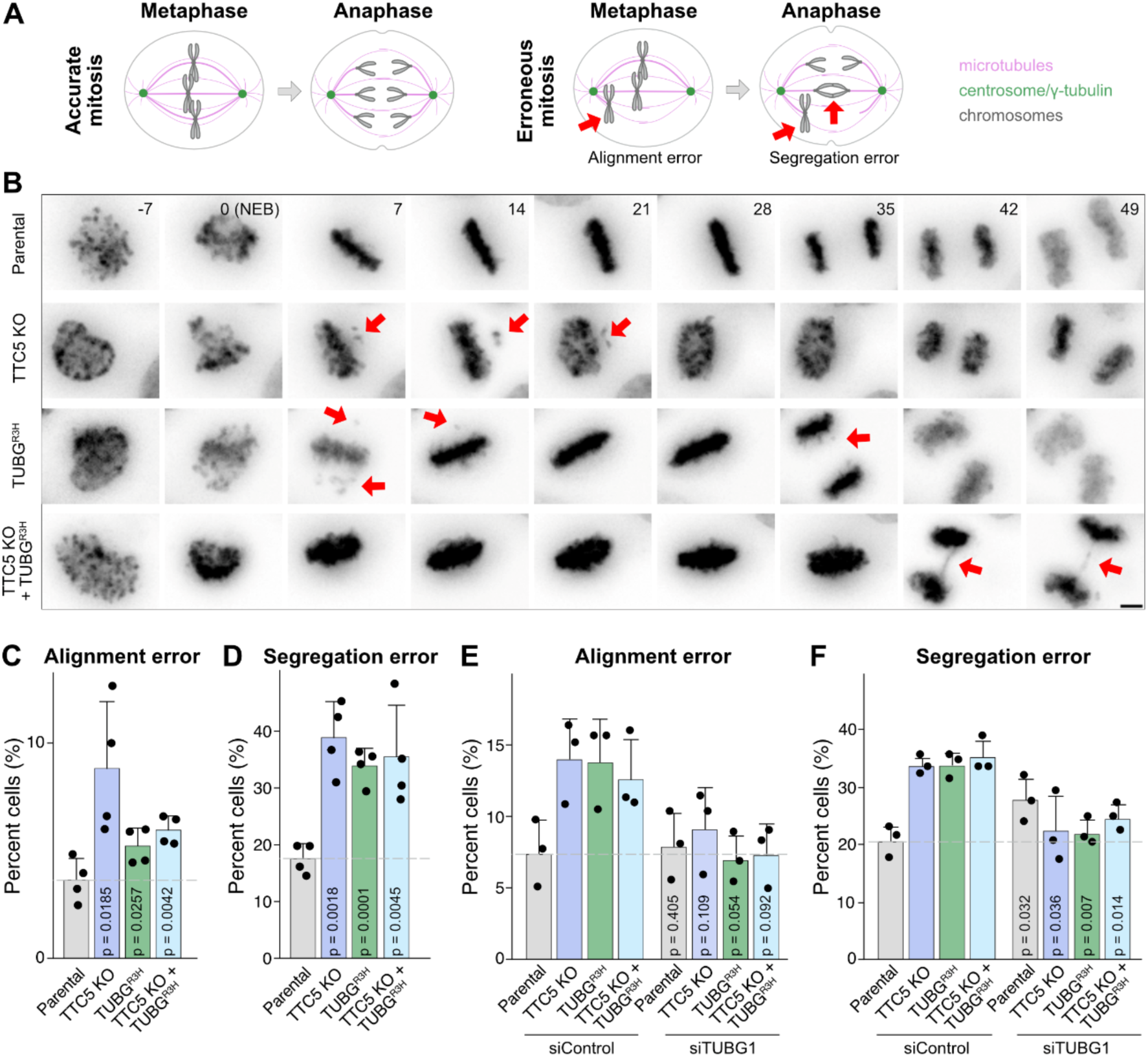
Regulated γ-tubulin biosynthesis maintains mitotic fidelity. (**A**) A schematic illustrating accurate and erroneous mitosis, with chromosome alignment errors during metaphase and chromosome segregation errors in anaphase. (**B**) Representative live-cell images of HeLa parental, TTC5 KO, TUBG^R3H^, and TTC5 KO + TUBG^R3H^ cell lines progressing through mitosis. DNA was visualized using Spy555-DNA dye, and maximum-intensity projections of 3D volumes are shown. Frames are aligned to nuclear envelope breakdown (NEB, t = 0). Chromosome alignment and segregation errors are indicated with red arrows. Scale bar = 10 μm. (**C-D**) Occurrence of chromosome alignment errors in metaphase (C) and chromosome segregation errors in anaphase (D) in the indicated cell lines. Data are presented as mean ± SD from four independent replicates, and 337, 340, 358, and 287 analysed Parental, TTC5 KO, TUBG^R3H^, TTC5 KO + TUBG^R3H^ cells, respectively. Indicated are *p*-values in unpaired, two-tailed Student’s *t*-tests for each of the cell lines with the parental cell line as reference. (**E-F**) Frequency of chromosome alignment (E) and chromosome segregation errors (D) in control (siControl) and TUBG1-depleted (siTUBG1) HeLa parental, TTC5 KO, TUBG^R3H^, and TTC5 KO + TUBG^R3H^ cells. Data are presented as mean ± SD from three independent replicates and over 100 analyzed cells per genotype (Parental: 166 siControl and 177 siTUBG1 cells; TTC5 knockout: 156 siControl and 181 siTUBG cells; TUBG^R3H^: 184 siControl and 129 siTUBG cells), and TTC5 knockout + TUBG^R3H^: 168 siControl and 182 siTUBG cells)). Indicated are Holm-adjusted *p*-values in unpaired, two-tailed Student’s *t*-tests for each genotype, with the corresponding siControl genotype as a reference.

We hypothesized that the mitotic defects observed in TTC5 KO, TUBG^R3H^, and TTC5 KO + TUBG^R3H^ cells might arise from the elevated γ-tubulin levels in these lines. To test this, we established a condition where γ-tubulin was acutely depleted by ∼20-30% using siRNA (**Fig. S7B-C**) and then examined mitotic fidelity across the different mutant cell lines. In line with our hypothesis, partial depletion of γ-tubulin restored mitotic fidelity to levels comparable to those of control cells (**Fig. 5E-F**), consistent with a causal role for γ-tubulin abundance.

Overall, these findings demonstrate that γ-tubulin mRNA regulation ensures mitotic fidelity by maintaining appropriate γ-tubulin levels.

## Discussion

Historically, selective tubulin mRNA degradation via tubulin autoregulation has been examined exclusively in the context of α- and β-tubulin-encoding transcripts, long before γ-tubulin was identified [20,21,46]. Recent mechanistic studies have revealed that activation of the tubulin autoregulation pathway triggers the release of TTC5 from soluble αβ-tubulins, enabling it to engage nascent α- and β-tubulin chains on translating ribosomes [22,24]. Through recruitment of SCAPER and the CCR4-NOT complex, TTC5 promotes mRNA deadenylation and decay [25]. This pathway was thought to exclusively regulate α- and β-tubulin mRNAs.

In this study, we identify γ-tubulin as a previously unrecognized target of the tubulin autoregulation pathway and show that its mRNA is degraded through the TTC5–SCAPER–CCR4-NOT axis in response to elevated soluble αβ-tubulins (**Fig. 2**). Loss of this regulatory mechanism, either through TTC5 knockout or mutagenesis of the γ-tubulin autoregulatory motif MPREI (γ-tubulin^R3H^ mutant), results in a modest yet reproducible increase in γ-tubulin protein levels (**Fig. 3**). In turn, this elevates centrosomal γ-tubulin during mitosis (**Fig. 3**), enhances microtubule nucleation capacity (**Fig. 4**), and reduces mitotic fidelity (**Fig. 5**).

Our identification of γ-tubulin as a target of the same TTC5–SCAPER–CCR4-NOT pathway that regulates α- and β-tubulin reveals tubulin autoregulation as a coordinated system rather than parallel, class-specific mechanisms. By using a common molecular machinery to control all three tubulin classes, cells are positioned to synchronously adjust microtubule building blocks and nucleation capacity in response to changes in soluble tubulin pools. Such coordination is likely essential to maintain an appropriate balance between microtubule mass, number, and organization, ensuring that heterodimer availability and nucleation output scale together rather than independently. This view extends tubulin autoregulation from a mechanism that buffers αβ-tubulin excess to a system-level regulator of the microtubule network.

Intriguingly, among γ-TuRC components, only γ-tubulin mRNA is subject to autoregulation in response to microtubule destabilization. While monomeric γ-tubulin can nucleate microtubules *in vitro* [47,48], whether this is functionally relevant *in vivo* remains unclear, since γ-TuRCs serve as the predominant nucleation templates under physiological conditions. This selective control raises the question as to why cells regulate a single subunit of such a large multiprotein complex. A possible explanation lies in the central role of γ-tubulin in stabilizing other γ-TuRC components. Previous work has shown that γ-tubulin, GCP2, and GCP3 are co-stabilized through their assembly into γ-tubulin small complexes (γ-TuSCs) [49]. Consistent with this, several identified ubiquitination sites in GCP2 and GCP3 are located in their γ-tubulin–binding regions, suggesting that unbound GCPs are selectively degraded [49–53]. Because γ-tubulin also forms γ-TuSC–like assemblies with GCP4–6, its abundance likely influences the stability of the broader γ-TuRC complex. Thus, regulating γ-tubulin alone may provide cells with an efficient way to tune γ-TuRCs abundance.

Tubulin autoregulation has previously been shown to be important for maintaining mitotic fidelity, yet the mechanism underlying this process remained unclear [22,24,25]. Our findings show that tubulin autoregulation-associated mitotic defects are largely driven by elevated γ-tubulin protein levels (**Fig. 5E-F**). This is evidenced by the mimicry in mitotic defects in TTC5 KO and γ-tubulin^R3H^ mutant cells. Further supporting this view, subtle γ-tubulin depletion restored mitotic fidelity to the levels seen in parental cells. This finding may seem surprising, given that mitotic defects arising from disrupted tubulin autoregulation would more intuitively be attributed to changes in α- and β-tubulin, as previously proposed [54]. Alterations in γ-tubulin quantity, however, have long been associated with defects in mitotic spindle organization [9,28–31,55,56]. While most previous studies have relied on overexpression or knockdown approaches that cause drastic perturbations in protein abundance, we demonstrate that even subtle changes in γ-tubulin levels are sufficient to compromise mitotic fidelity. Our work highlights γ-tubulin–mediated nucleation as a particularly sensitive control point for mitosis and suggests that tubulin autoregulation safeguards mitotic fidelity primarily by constraining nucleation output.

A surprising outcome was that the γ-tubulin^R3H^ point mutant and TTC5 KO did not show additive defects in mitotic errors, despite each perturbation increasing segregation errors when tested alone. Compared to TTC5 KO alone, the R3H allele in the TTC5 KO background produced a small but detectable increase in centrosomal nucleation output. γ-TuRC activity is exquisitely sensitive to sequence alterations because the complex is large and highly cooperative, so many mutations produce outsized functional consequences [57–59]. However, the R3H change appears to be sufficiently mild that it does not further exacerbate the mitotic defects caused by loss of TTC5.

Several perturbations of γ-tubulin or centrosome function result in abnormal mitotic spindles and produce moderate increases in chromosome missegregation (on the order of tens of percent) rather than wholesale spindle collapse [55,60–62]. For example, mutations of γ-tubulin in *Schizosaccharomyces pombe* lead to increased DNA missegregation events ranging from 3.33% to 27.04% [60]. Meanwhile, in DT40 chicken cells, knockout of CEP152 (Centrosomal Protein 152) or STIL (SCL/TAL1 interrupting locus)—proteins that are essential to centriole formation—leads to ∼30% rate in chromosome missegregation events [62], illustrating that mitotic fidelity can tolerate some change in nucleation or spindle pole integrity before complete failure occurs.

Recently, TTC5 and SCAPER homologs in *Caenorhabditis elegans* (*C. elegans*) were observed to control neuronal microtubule network architecture by recruiting γ-tubulin and promoting γ-tubulin-mediated nucleation, independently of tubulin autoregulation [63]. Whether this autoregulation-independent function of TTC5, SCAPER, and γ-tubulin is conserved across different cell types and in humans is unknown. However, neither γ-tubulin nor any other γ-TuRC components were identified as interactors of either TTC5 or SCAPER in previous proximity-labeling experiments [22,25], suggesting that, at least in cultured cells, this interaction likely does not occur. Interestingly, unlike α- and β-tubulin, whose amino-terminal autoregulatory motifs are conserved in worms, the amino-terminal motif of γ-tubulin differs drastically in *C. elegans* (MSGTG) compared to that of humans and lacks the critical amino acids R3 and E4 required for autoregulation. This raises the intriguing possibility that TTC5 and SCAPER may have co-evolved in worms to regulate γ-tubulin function through direct recruitment, whereas in humans, TTC5 and SCAPER modulate γ-tubulin function by influencing its protein abundance.

In summary, our study expands the scope of tubulin autoregulation beyond α- and β-tubulins, identifying γ-tubulin as a quantitatively regulated target whose abundance fine-tunes microtubule nucleation and mitotic accuracy. These findings uncover a previously unappreciated layer of post-transcriptional control governing microtubule-nucleating activity and highlight how subtle changes in γ-tubulin levels can alter genome inheritance. Because chromosome missegregation can lead to cell cycle arrest, genomic instability, abnormal karyotypes, and ultimately tumorigenesis or cell death, understanding the sources of compromised mitotic fidelity is of key interest [64]. Given the widespread upregulation of γ-tubulin in tumours, the TTC5-dependent pathway described here offers a new conceptual and mechanistic entry point for exploring how γ-tubulin quantity contributes to pathological microtubule organization in cancer and developmental disease [32, 65–70].

## Materials and methods

### Cell culture, transfection, and treatment

Flp-In T-Rex HeLa cells (Thermo Fisher Scientific, R71407) and Flp-In T-Rex HEK293 cells (Thermo Fisher Scientific, R78007) were maintained in DMEM with GlutaMAX (Thermo Fisher Scientific, 10566016) supplemented with 10 % heat-inactivated Fetal Bovine Serum (FBS, Pan Biotech, P30-3306) and 1 % vol/vol penicillin-streptomycin (Thermo Fisher Scientific, 15140122) at 37 **°**C in a humidified atmosphere containing 5 % CO_2_. hTERT RPE1 FRT/TR (a kind gift from the Jonathon Pines lab) cells were cultured in DMEM/F12 (Thermo Fisher Scientific, 11320033) supplemented with 10 % FBS and 1 % vol/vol penicillin-streptomycin at 37 **°**C in a humidified atmosphere containing 5 % CO_2_.

Stable cell lines expressing wild-type TTC5, autoregulation-defective TTC5^R147A^ mutant, wild-type SCAPER, autoregulation-defective SCAPER^Δ620^, and wild-type CNOT11 were generated previously [22,23]. Cell lines with stable expression of TTC5^DE-AA^ mutant, γ-tubulin-eGFP, γ-tubulin-eGFP ^R3H^-eGFP, γ-tubulin-FLAG or γ-tubulin^R3H^-FLAG were generated from the Flp-In HeLa cells according to the manufacturer’s protocol (K650001). Briefly, transgenes were subcloned into a pcDNA5/FRT vector and co-transfected with pOG44 (Flp recombinase) at a 1:1 ratio using Lipofectamine 3000 (Thermo Fisher Scientific, L300015). The following day, cells were selected with hygromycin B (0.2 mg/ml; Corning, 30-240-CR) and blasticidine S (10 µg/mL; Thermo Fisher Scientific, R21001) for 10-15 days. Transgene expression was induced with doxycycline (400 ng/ml) for 24 hours.

Colchicine, nocodazole, combretastatin A4, and paclitaxel treatments were performed in complete growth media for the indicated durations at 1 µM concentrations unless stated otherwise. For RNA interference, cells were transfected with TUBG1 siRNA (1 hs.Ri.TUBG1.13.1; Integrated DNA Technologies) or a control non-targeting siRNA (51-01-14, Integrated DNA Technologies) using Lipofectamine RNAiMAX (Invitrogen, 13778075), according to the manufacturer’s instructions.

### CRISPR-Cas9 Editing of cells

CRISPR-Cas9 editing strategies were designed using the IDT HDR Design tool. To generate *TUBG1^R3H^* and *TUBG2^R3H^* mutant cell lines, *TUBG1* was edited first, followed by *TUBG2* in HeLa Flp-In TRex or HeLa Flp-In TRex TTC5 knockout cells.

To generate TUBG1^R3H^ mutant cells, cells were transiently transfected with Alt-R™ CRISPR-Cas9 gRNA targeting *TUBG1* (Integrated DNA Technologies, Hs.HC9.DSGX4159.AA or Hs.HC.DSGX4157.AA), Alt-R HDR Donor Oligo (Integrated DNA Technologies, CD.HC9.QZNR1251+), and Alt-R S.p. Cas9-GFP V3 (Integrated DNA Technologies, 10008100 Lot n°914743) using Lipofectamine RNAiMAX (Thermo Fisher Scientific, P13778075) following the manufacturer’s protocols. Alt-R HDR Enhancer V2 (Integrated DNA Technologies, 10007910 Lot n°914418) was added to promote homology-directed repair as per the manufacturer’s instructions.

30 hours post-transfection, GFP-positive cells were sorted using Cell Sorter SONY SH800 (Access, Geneva) and seeded into 96-well plates at a density of 1 cell per well to obtain single clones. Correctly edited clones were identified by PCR amplification of the targeted region, followed by Sanger sequencing, which was analyzed using CLC Genomics Workbench 24.

HeLa Flp-In TRex TUBG1^R3H^ or HeLa Flp-In TRex TTC5 knockout + TUBG1^R3H^ cells were then subjected to a second round of CRISPR-Cas9 editing by transiently transfecting them with Alt-R™ CRISPR-Cas9 gRNA targeting *TUBG2* (Integrated DNA Technologies Hs.HC9.DSGY1116.AA), Alt-R HDR Donor Oligo (Integrated DNA Technologies, CD.HC9.HVPJ2280+), and Alt-R S.p. Cas9-GFP V3, following the same transfection and selection procedures described above. Correctly edited clones were identified by PCR amplification of both targeted regions, followed by Sanger sequencing and analysis using CLC Genomics Workbench 24.

### Profiling RNA levels via RT-qPCR

Cells were grown to ∼70% confluence in 100 mm dishes and treated with DMSO (vehicle control) or microtubule poisons. Cells were harvested by scraping in RA1 lysis buffer and total RNA was isolated using the NucleoSpin RNA Mini Kit for RNA Isolation (Macherey-Nagel, 740955) according to the manufacturer’s protocol. Following RNA extraction, 1 μg of RNA was used to generate cDNA with SuperScript^TM^ IV First-Strand Synthesis System Kit (Thermo Fisher Scientific 18091050) and random hexamer primers following the manufacturer’s instructions. RT-qPCR was carried out using 10 ng of cDNA and 2× PowerUp SYBR Green master mix (Life Technologies, A25777) and the indicated primers on a BioRad thermocycler (BioRad). Relative expression was analyzed using the ddCt method [71]. All data were normalized to housekeeping reference genes and to DMSO-treated controls. Experiments include at least three biological replicates. Statistical analysis and data plotting were performed using R/Rstudio (version 4.4.1). All primers used for qPCR are listed below.

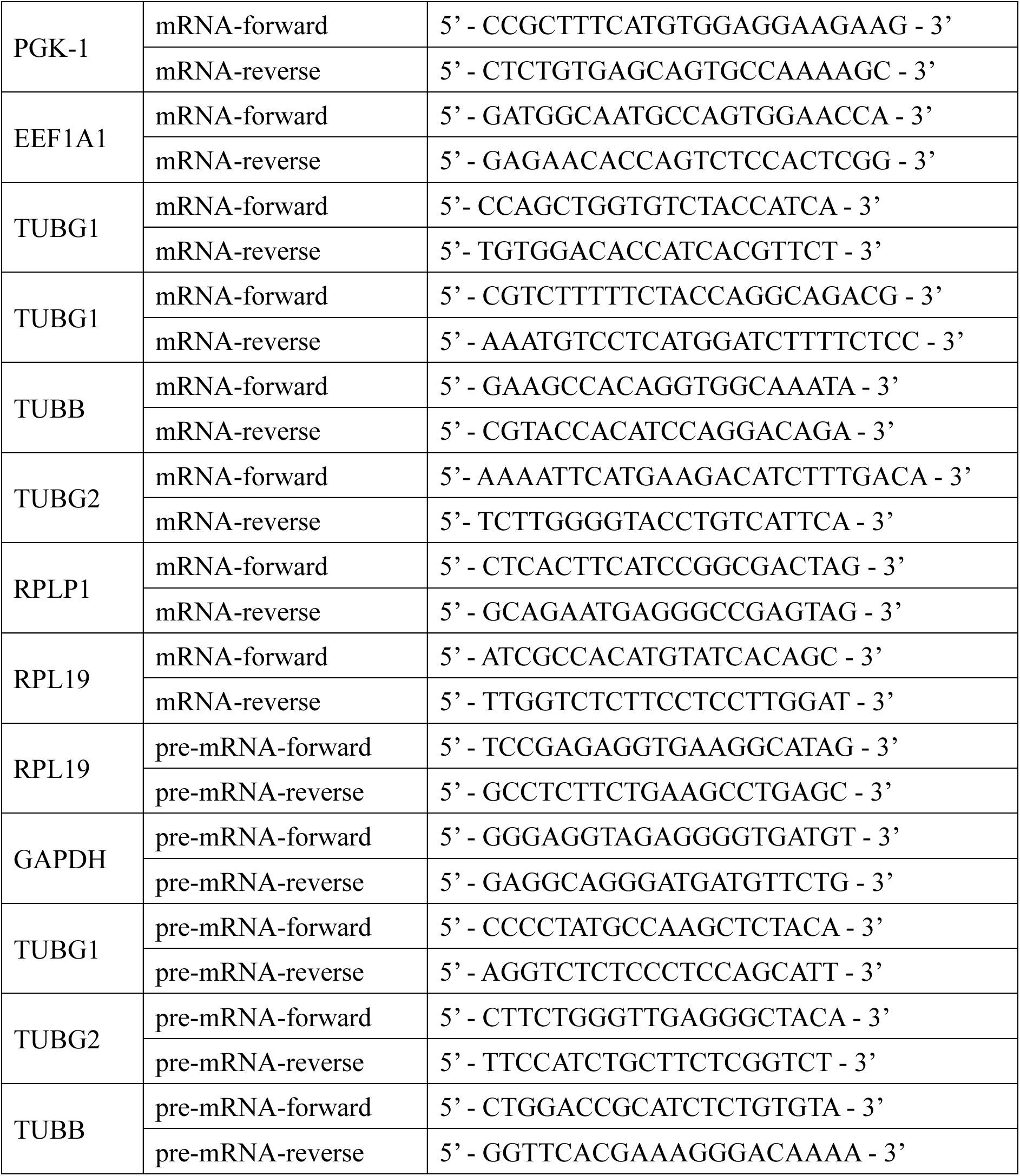

### Recombinant protein purification

To express 6XHis-StrepII tagged wild-type TTC5, pET28a plasmids containing sequence-verified vectors were transformed into *E. coli* expression strains BL21 by the heat shock method. Starter cultures were prepared by inoculating cultures from frozen glycerol stocks in 100 mL Lysogeny Broth (LB) medium containing kanamycin 50 μg/mL at 37 °C overnight in a shaking incubator. The following day, starter cultures were diluted 1:20 in 1 liter LB kanamycin and grown in a shaking incubator at 37 °C until reaching an OD of 0.6 to 0.8. At this point, protein expression was induced by the addition of 0.2 mM IPTG at an A600 of 0.6 at 18 °C in a shaking incubator overnight. Bacterial cells were harvested by centrifugation and pellets were resuspended in 50 mL chilled lysis buffer [50 mM HEPES pH 7.4, 500 mM NaCl, 20 mM imidazole, 1 mM PMSF, 1 mM TCEP] before being lysed by passing five times through the French press at 1,000 psi. The lysate was then centrifuged at 20,000 g and consequently loaded onto a 1 mL HisTrap column (HISPUR Ni-Nta Spin columns, Thermofisher) pre-equilibrated in lysis buffer. Columns were washed with ∼ 10 column volumes of lysis buffer and eluted in lysis buffer containing 250 mM imidazole. The elute was then loaded onto a Strep-Tactin column (StrepTrap XT prepacked chromatography column, Cytiva) prewashed with 5 mL pure water and 5 mL Strep Buffer [20 mM HEPES pH 7.4, 500 mM NaCl, 1 mM TCEP]. After washing the column with 10 column volumes of Strep Buffer, Strep Buffer containing 50 mM biotin was used to elute bound TTC5, which was then dialyzed against Strep Buffer and concentrated using an Amicon Ultra 4 centrifugal filter unit (Millipore). Purified protein was verified using a western blot, flash frozen, and stored at-80 °C until use.

### Recombinant TTC5 Pulldown of Cell Lysates

TTC5 knockout cells were grown to 80% confluence in a 145 mm dish and treated with DMSO control or combretastatin A4 (10 μM) for 1 hour. Cells were collected after centrifugation at 350 g and lysed with lysis buffer [50 mM HEPES pH 7.4, 100 mM KAc, 5 mM MgAc2, 1mM DTT, 100 μg/mL digitonin, 1 x EDTA-free protease inhibitor cocktail (Roche)] for 20 minutes on ice. Lysates were then cleared by centrifugation at 20,000 g for 15 minutes at 4 °C. A small fraction of lysates was used for total RNA extraction and analyzed for tubulin mRNAs by RT-qPCR as described above. The remainder of the lysates was incubated with 500 nM recombinant TTC5 for 4 minutes before incubation with 15 μL Streptactin Sepharose Beads (StrepTactin Sepharose High Performance, Cytiva) for 2 hours at 4 °C with head-to-tail rotation to recover TTC5 and all bound components. Beads were washed four times with 400 μL PSB [50 mM HEPES pH 7.4, 100 mM KAc, 2 mM MgCl2] and eluted with 50 μL 50 mM biotin in PSB at 4 °C with head-to-tail rotation for 60 minutes. An aliquot of eluted samples was analyzed by SDS-PAGE and SYPRO Ruby stain to visualize TTC5 and associated proteins. Another aliquot of the eluted products was used to isolate RNA, generate cDNA, and perform qPCR as mentioned previously. Data analysis was performed using the ddCt method [43]. Data obtained for genes of interest were normalized to the housekeeping gene RPLP1 and DMSO-treated cells.

### Western Blotting and Quantification

Indicated cell lines were lysed in RIPA buffer [50 mM Tris-HCl, pH 8.0, 150 mM NaCl, 1% Triton X-100, 0.5% sodium deoxycholate, 0.1% SDS] supplemented with protease (1:100, Halt^TM^ Thermo Fisher Scientific 78420) and phosphatase inhibitors (1:25, PhosSTOP, Roche 4906845001). Lysate concentrations were determined by Pierce BCA assay kit (Thermo Fisher Scientific).

Lysates were separated using SDS-PAGE on 4-12 %, 8% or 10 % Bolt^TM^ Bis-Tris Plus Mini Protein Gels (Thermo Fisher Scientific NW04125BOX; NW00080BOX; NW00105BOX), and transferred to 0.2 μm nitrocellulose membranes (Amersham, Cytivia). Membranes were blocked in 5 % non-fat dry milk prepared in PBS containing 0.2 % Tween-20. Primary antibodies anti-γ-tubulin (1:2000, Sigma-Aldrich T3320) and anti-GAPDH (1:5000, Thermo Fisher Scientific MA515738) were incubated for 1 h at room temperature or 4 °C overnight. Detection was done using IRDye-conjugated secondary antibodies LI-COR 680 (1:10000, Thermo Fisher Scientific A32729) and LI-COR 800 (1:10000, Thermo Fisher Scientific, A32735). Band intensities were quantified in Fiji (ImageJ, 1.54f). Briefly, rectangular regions of interest were manually drawn around each band, keeping the area constant throughout all lanes. Upon background subtraction, mean intensity values of γ-tubulin were normalized to the corresponding GAPDH signals. Data were expressed as γ-tubulin/GAPDH ratios, and technical replicates averaged before plotting and statistical analyses.

For Flag-co-immunoprecipitation experiments, membranes were incubated with anti-GCP4 (1:100, Santa Cruz Biotechnology sc-271876), anti-GCP2 (1;1000, Proteintech 25856-1-AP), anti-GCP3 (1:750, Proteintech 15719-1-AP), and anti-Flag (1:2000, Sigma-Aldrich F3165-1MG). For recombinant TTC5 pulldown lysates, membranes were incubated with anti-α-tubulin antibody (1:10000, Thermo Fisher Scientific 14-4502-37), anti-Strep antibody (1:2000 Abcam ab76949), 1:500 anti-RPL8 antibody (1:500 Abcam ab169538).

### Immunofluorescence

Cultured cells plated on presterilized glass coverslips were fixed in cold methanol (-20 °C) for 10 minutes. Following 3x washes with PBS, the cells were incubated at room temperature for 1 hour with blocking solution 2 % BSA diluted in PBS with 0.02 % Tween-20 (PBS-T). The samples were then treated with primary antibodies such as anti-γ-tubulin (1:500, Sigma-Aldrich T3320), anti-α-tubulin (1:2000, Thermo Fisher Scientific 14-4502-37), and anti-centrin (1:500, Merck Millipore 04-1624) diluted in blocking solution and incubated at room temperature for 1 hour. Following 3x washes with PBS-T, cells were incubated for 1 hour at room temperature with the following secondary antibodies: Alexa 488 (1:1000, Thermo Fisher Scientific, A32731, Lot n° WC318798) and 555 (1:1000, Thermo Fisher Scientific, A32727, Lot n° WA316324), together with DAPI (1:5000, 5mg/mL, Thermo Fisher Scientific, D1306). After 3x washes with PBS-T, the cells were mounted onto glass slides with mounting medium [20mM Tris, pH 8.0, 0.5% N-propyl gallate, 50-90% Glycerol]. Images were taken on a widefield Nikon Eclipse Ti2-E inverted microscope using NIS Elements (Nikon) and 60× Plan Apochromat Lambda oil objective (NA 1.4, Nikon). To quantify γ-tubulin/centrin ratios, a circular region of interest (ROI) was drawn around the centrosome as indicated by γ-tubulin and centrin. Fluorescence measurements were obtained from the ROI spanning eight consecutive Z-stacks (2.4 μM total) centered on the centrosomal plane. The ROI size was kept constant throughout all samples and channels. Following background subtraction, integrated density measurements were taken.

### Microtubule Regrowth Assay

Cells were grown on presterilized coverslips in 6-well plates and incubated in 4 °C growth medium on ice for 1 hour to depolymerize microtubules. Coverslips were then washed once with 4 °C cold PBS and incubated with pre-warmed growth medium (37 °C) for 45 seconds to allow microtubule regrowth, after which cells were fixed with cold methanol. Cells were then processed for immunofluorescence as described above using the following primary antibodies: as anti-γ-tubulin 1:500 (Sigma-Aldrich T3320), anti-α-tubulin 1:2000 (Thermo Fisher Scientific 14-4502-37). Only metaphase cells were imaged and analyzed. Z-stacks were acquired at 0.3 μM intervals, spanning a total thickness of 12 μM using NIS Elements (Nikon) and 60× Plan Apochromat Lambda oil objective (NA 1.4, Nikon). Image analysis was performed using ImageJ. Specifically, fluorescence measurements were obtained from the sum-projection of six consecutive Z-slices centered on the centrosomal plane. A circular ROI was drawn around the centrosome as indicated by γ-tubulin staining. The ROI size was kept constant throughout all samples and channels. Following background subtraction, integrated density measurements were taken. Data are presented as α-tubulin fluorescent intensity or as α-tubulin/γ-tubulin ratios.

### Live cell imaging and data analysis

Flp-In T-REx HeLa cells of the genotypes indicated in the figure legends were plated in 8-well 1.5 Ibidi glass coverslip (Ibidi 80807) in regular growth medium. Following 24 hours, the medium was changed to DMEM, high glucose, HEPES without phenol-red (Thermo Fisher Scientific, 21063-029) supplemented with 10 % FBS, 1% vol/vol penicillin-streptomycin, and 1:1000 SPY650-DNA (Spirochrome, SC501) for 2 hours or 1:1000 SPY555-DNA (Spirochrome, SC201) for 6 hours prior to imaging. Time-lapse images were acquired using a Nikon Eclipse Ti2-E inverted equipped with a perfect focus system and an incubation chamber at 37 °C with controlled humidity and 5 % CO2 (OkoLab). Three-dimensional images at multiple stage positions were acquired in steps of 2 μm, every 7 minutes for 3.5 to 5 hours using NIS Elements (Nikon) and 20× Plan Apochromat Lambda objective (NA 0.80, Nikon). Mitotic cells were analyzed using 3D reconstructions in Fiji. Unaligned chromosomes in metaphase and chromosome segregation errors in anaphase were scored based on the Spy650-DNA or SiR-DNA signal. At least 100 cells per cell line in three or more biological replicates were documented using Excel and processed and plotted using R (version 4.4.1). Statistical analyses were performed in R. Chromosome alignment errors were classified as events where not all the chromosomes were properly aligned on the spindle equator in metaphase and/or anaphase. Meanwhile, chromosome segregation errors were classified as instances where sister chromatids failed to properly separate, either by segregating both in the same daughter cell or by forming a bridge in anaphase. Numbers reported represent the percentage of cells experiencing abnormality. To generate representative examples of mitosis, maximum intensity projections and inverted color profiles of examples were prepared in Fiji (ImageJ, version 2.14.0/1.54f) and exported as still images.

### Microarray data analysis

Publicly available microarray datasets were read, normalized, and analyzed using linear models and empirical Bayes methods for differential expression in the limma package [72]. Microarray probes annotated to the TUBG1 and TUBG2 isoforms were used for analysis. Heatmaps of differential expression were generated using R.

## Statistical analysis

For all data presented in figures, the number of independent biological replicates, statistical test with *p*-values, and, where applicable, the number of analyzed cells, are indicated in the figures and figure legends. No statistical method was used to predetermine the sample size. The experiments were not randomized, and the investigators were not blinded to allocation during experiments and outcome assessments. Statistical analyses were performed in R and GraphPad Prism 8.

## Acknowledgments

We are grateful to A. Almeida and A. Noireterre for the critical reading of the manuscript, E. Vartholomaiou for help with protein purification, and all members of the Gasic team for their valuable feedback and support. The authors thank Prof. Jonathon Pines, Prof. Paul Guichard, and Dr. Virginie Hamel for sharing cell lines and reagents, Maté Fülöp for his help with structural analyses, and the Access Facility at the University of Geneva for their help with cell sorting. This work was supported by the Swiss National Science Foundation (SNSF, grant PCEFP3_194312 to I.G.) and the Republic and Canton of Geneva, Switzerland (DIP, to I.G.). I.G. was the Dale F. Frey Breakthrough Scientist of the Damon Runyon Cancer Research Foundation (DRG:227916).

## Author contributions

M.A. performed all experiments and conducted the formal data analysis. C.L. contributed to western blotting. M.A. prepared the figures and wrote the initial draft of the manuscript. I.G. conceived the study, oversaw its design and implementation, provided supervision and funding, and edited the manuscript.

## Supplementary figures and figure legends

**Figure S1.**
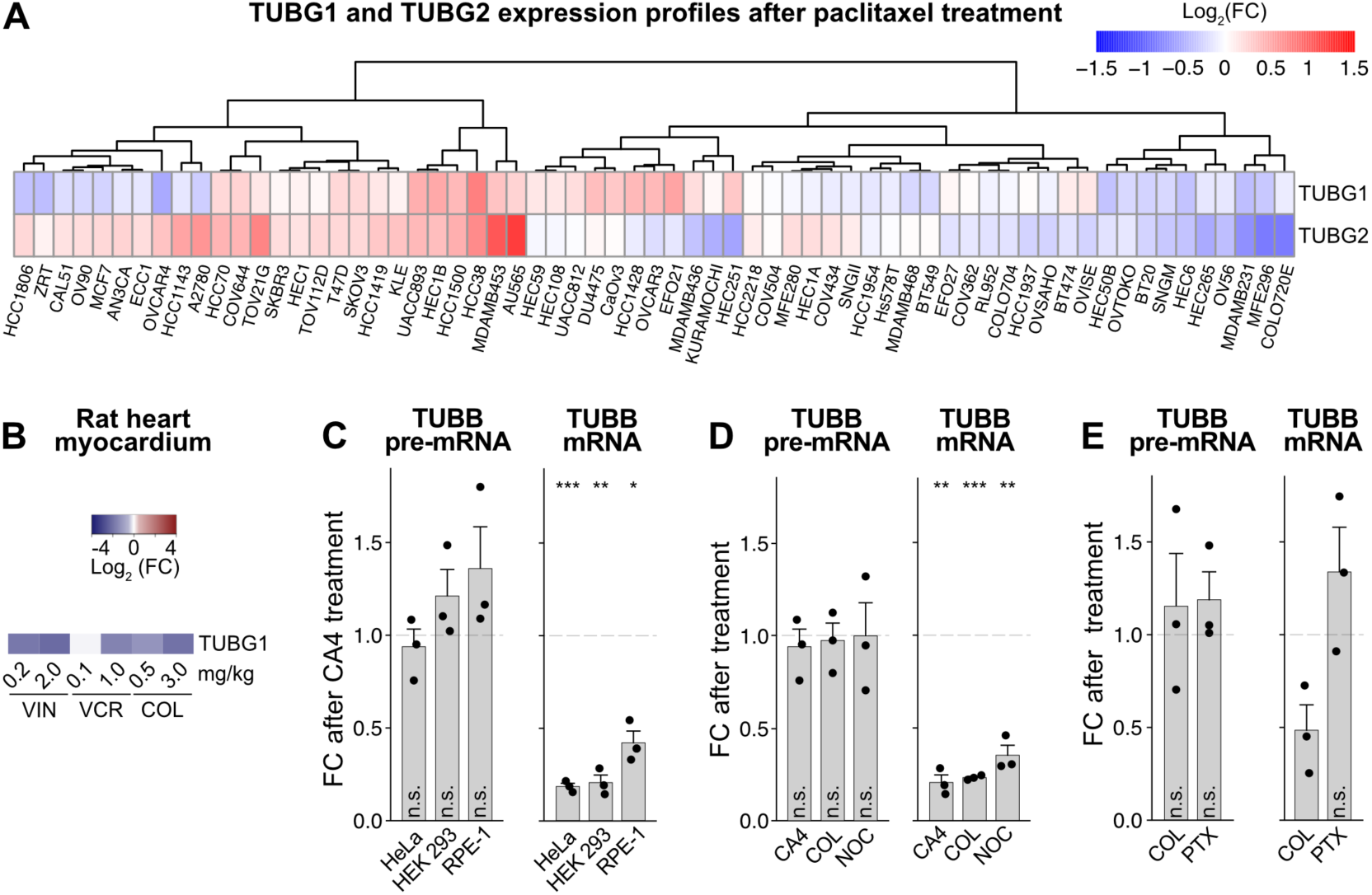
(related to Figure 1). (**A**) Relative expression levels of TUBG1 and TUBG2 in various cancer cell lines treated with paclitaxel for 24 h. Data were obtained from the GEO database (GEO series GSE50811, GSE50830, and GSE50831 [34]). Columns correspond to cells and rows to genes. Hierarchical clustering was performed on cells using Euclidean distance and complete linkage, and the results are displayed as a dendrogram above the heatmap, grouping cells with similar TUBG1/TUBG2 expression responses to treatment. Data are represented as Log_2_ Fold Change (Log_2_FC) relative to DMSO control, with the color key depicted on the right. (**B**) The color key and expression profiles of TUBG1 in rat heart endothelial cells isolated from control and the animals treated with microtubule poisons vinblastine (VIN), vincristine (VCR), or colchicine (COL) at indicated doses (x-axis) for 6 h. The data were retrieved from the GEO database (GSE19290). Data are represented as Log_2_Fold Change (Log_2_FC) relative to vehicle-treated control animals. (**C**) TUBB pre-mRNA and mRNA levels in HeLa, HEK293, and RPE1 cell lines following 6 hours of treatment with CA4. (**D**) TUBB pre-mRNA and mRNA levels in HEK293 cells after 6 hours of treatment with the indicated microtubule-targeting inhibitors. (**E**) TUBB pre-mRNA and mRNA levels in HeLa cells after 6 hours of treatment with 100 nM colchicine and 300 nM paclitaxel. All mRNA and pre-mRNA levels were normalized to the housekeeping transcripts RPL19 and GAPDH, respectively, and to DMSO-treated controls. Data represent mean ± SD from three independent experiments. A two-tailed Student’s *t*-test was performed for each of the cell lines with the DMSO-treated cells as reference. * *p* < 0.05, ** *p* < 0.01, *** *p* < 0.001, n.s. = not significant.

**Figure S2.**
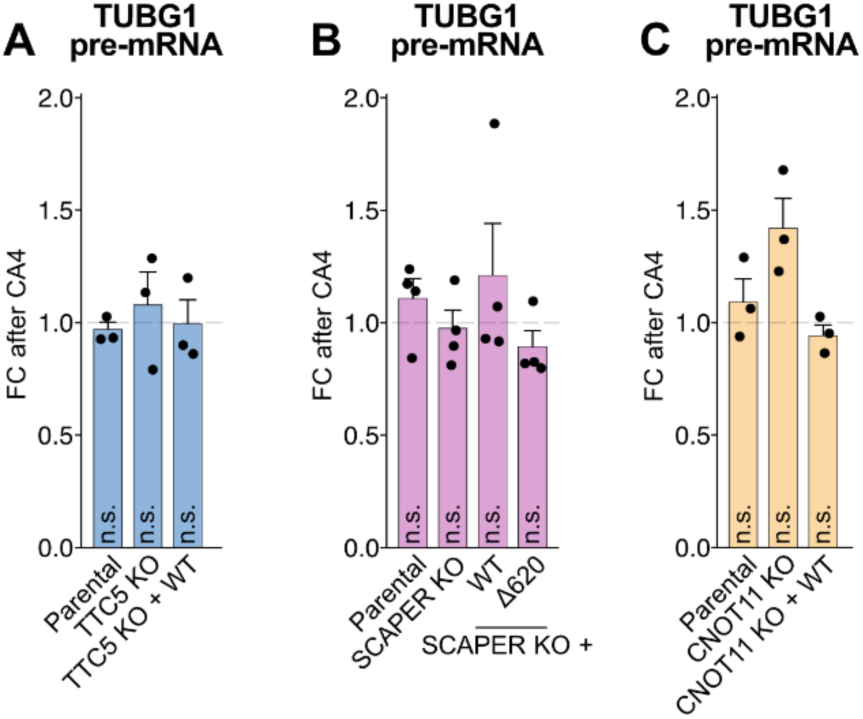
(related to Figure 2). (**A-C**) TUBG1 pre-mRNA levels following 8 hours of CA4 treatment in HeLa cells of the indicated genotypes. Data represent mean ± SD with each data point corresponding to an independent experiment (n=3 for panels A and C, and n=5 for panel B). Data were normalized to the housekeeping transcript GAPDH. A two-tailed Student’s *t*-test was performed for each of the cell lines with the DMSO-treated cells as reference. n.s. = not significant.

**Figure S3.**
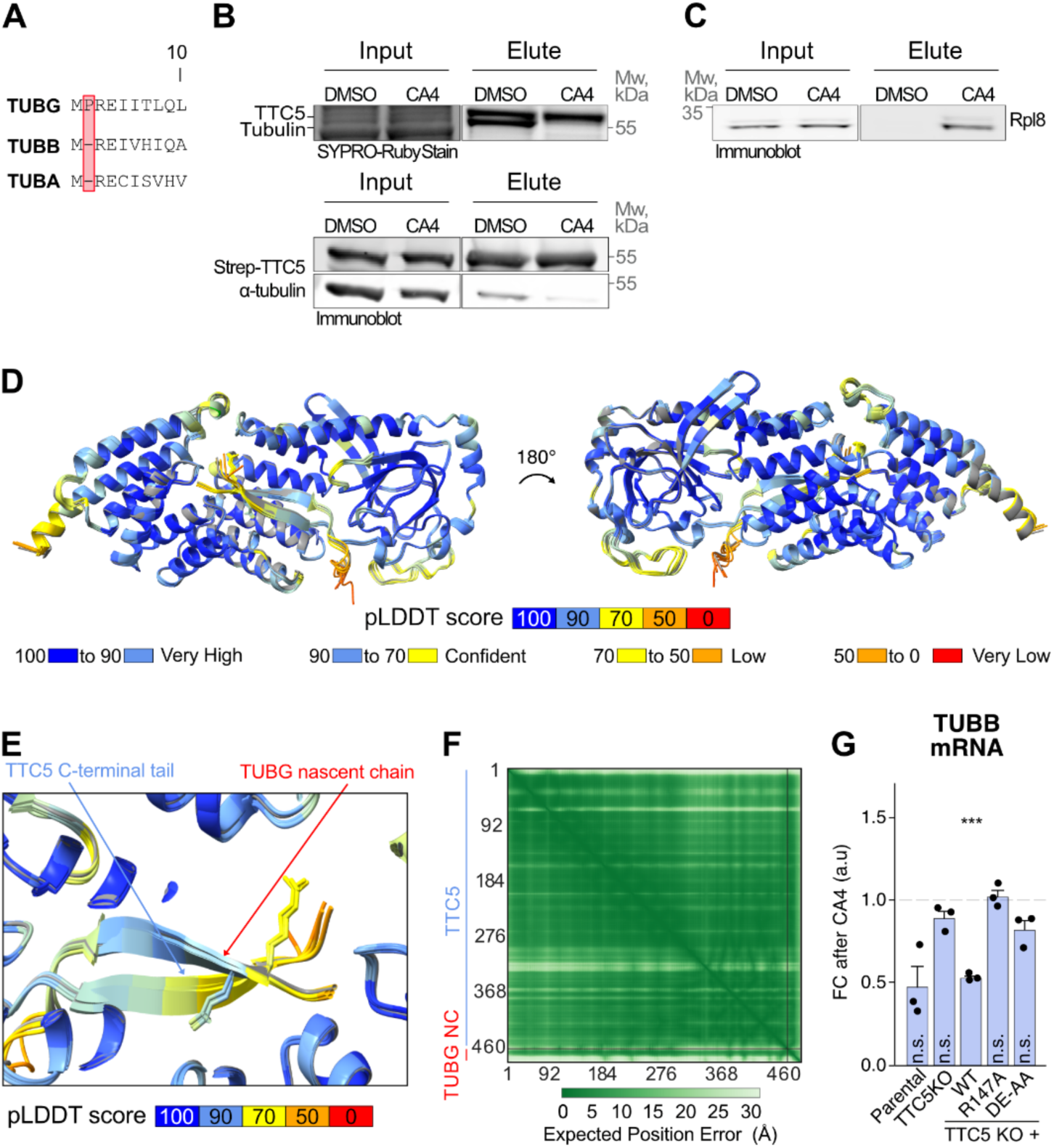
(related to Figure 2). (**A**) Sequence alignment of the amino-terminal motifs of human α-, β-, and γ-tubulins. (**B**) TTC5 KO HEK293 cells were pre-treated with DMSO (vehicle control) or combretastatin A4 for 1 hour, lysed, incubated with recombinant Strep-TTC5, and affinity-purified. Co-immunoprecipitated interactors were separated using SDS-PAGE, and Strep-TTC5 and α-tubulin were visualized using SYPRO-Ruby stain and western blot. (**C**) Interaction of ribosomal protein Rpl8 with Strep-TTC5 was visualized using a western blot. Experiments in (B) and (C) were repeated five times with similar results. (**D**) Superimposition of the five top AlphaFold3 predicted models of TTC5 with the first 20 amino acids of γ-tubulin. Residues are color-coded by predicted local distance difference test (pLDDT) scores, indicating the level of confidence in the prediction. The color scale is shown at the bottom. (**E**) Close-up view of the top five AlphaFold3 predicted models of TTC5 C-terminal tail and γ-tubulin nascent chain. The color key is depicted at the bottom. (**F**) Predicted aligned error (PAE) map of the AlphaFold3 model shown in Fig. 2G. The color scale is indicated at the bottom. (**G**) TUBB mRNA levels in TTC5 KO and indicated mutant TTC5 rescue cell lines following 8-hour treatment with CA4 (1 μM). Data represent mean ± SD of three independent replicates.

**Figure S4.**
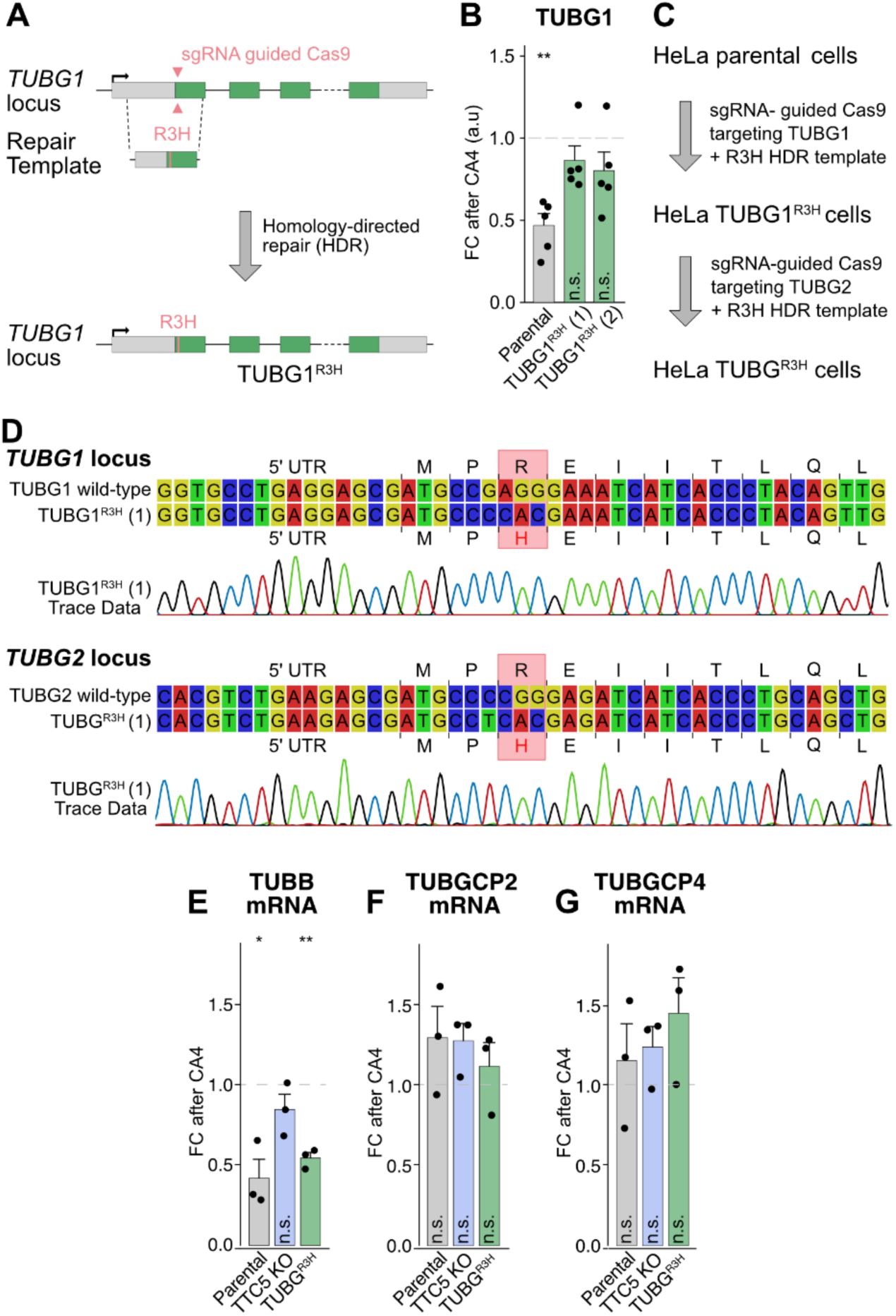
(related to Figure 2). (**A**) A schematic depicting the strategy used to generate *TUBG1^R3H^* mutant HeLa cells by CRISPR-Cas9 homology-directed repair. (**B**) TUBG1 mRNA levels in TUBG1^R3H^ cell lines in two independent clones (1 and 2) following CA4 treatment. Data represent mean ± SD from five independent replicates. (**C**) Scheme outlining the sequential CRISPR-Cas9 editing steps used to generate HeLa TUBG^R3H^ cells harboring both TUBG1^R3H^ and TUBG2^R3H^. A detailed description of the genome-editing strategy is provided in the Materials and Methods section. (**D**) Sanger sequencing results, including trace data visualized using CLC Genomics Workbench 24, highlighting the nucleotide substitutions introduced to generate the R3H mutation. In addition to the missense mutation, a synonymous mutation of the G/C nucleotide within the codon encoding Proline was introduced to prevent Cas9 re-cutting following homology-directed repair. (**E**) TUBB mRNA levels in TTC5 KO and indicated mutant TTC5 rescue cell lines following 8-hour treatment with 1 μM CA4, Data represents mean ± SD of three independent replicates. (**F-G**) TUBGCP2 (G) and TUBGCP4 (H) mRNA levels in parental, TTC5 KO, and TUBG^R3H^ cells following CA4 treatment. Data represent mean ± SD from three independent replicates, normalized to housekeeping transcript EEF1A1. A two-tailed Student’s *t*-test was performed for each of the cell lines with the DMSO-treated cells as reference. * *p* < 0.05, ** *p* < 0.01, *** *p* < 0.001, n.s. = not significant.

**Figure S5.**
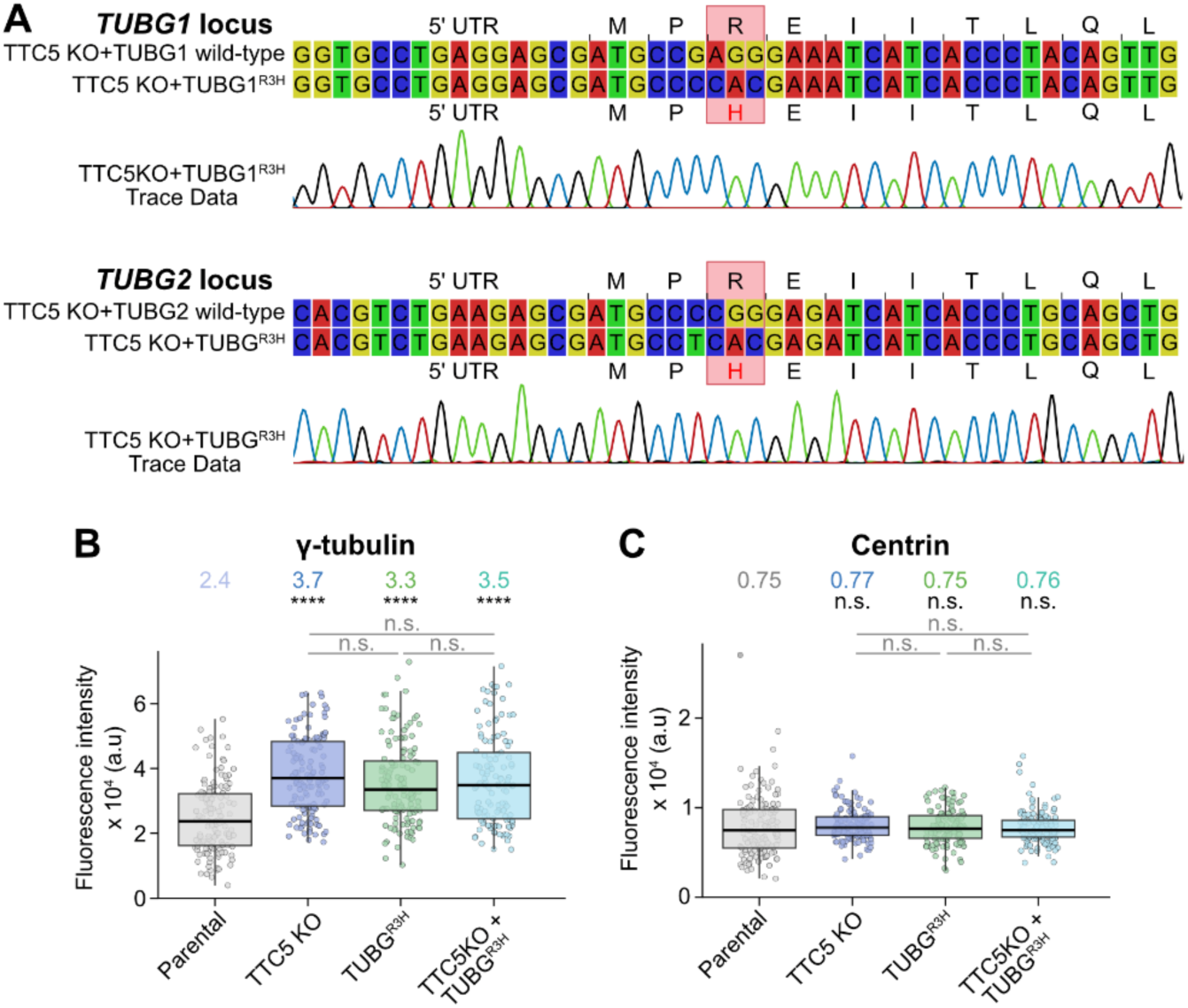
(related to Figure 3). (**A**) Sanger sequencing results of TTC5 KO + TUBG^R3H^ cells, including trace data visualized using CLC Genomics Workbench 24, highlighting the nucleotide substitutions introduced to generate the R3H mutation. In addition to the missense mutation, a synonymous mutation of the G/C nucleotide within the codon encoding Proline was introduced to prevent Cas9 re-cutting following homology-directed repair. (**B-C**) Quantification of centrosomal γ-tubulin (B) and centrin (C) fluorescence intensity analysed in Fig. 3D. Data show median ± interquartile range. Median values colour-coded by genotype are indicated above the respective boxplots. Shown are Benjamini-Hochberg-adjusted *p-*values from an unpaired Mann-Whitney test comparing ranks for each of the indicated cell lines with the parental cell line as reference, or across the groups. * *p* < 0.05, ** *p* < 0.01, *** *p* < 0.001, **** *p* < 0.0001, n.s. = not significant.

**Figure S6.**
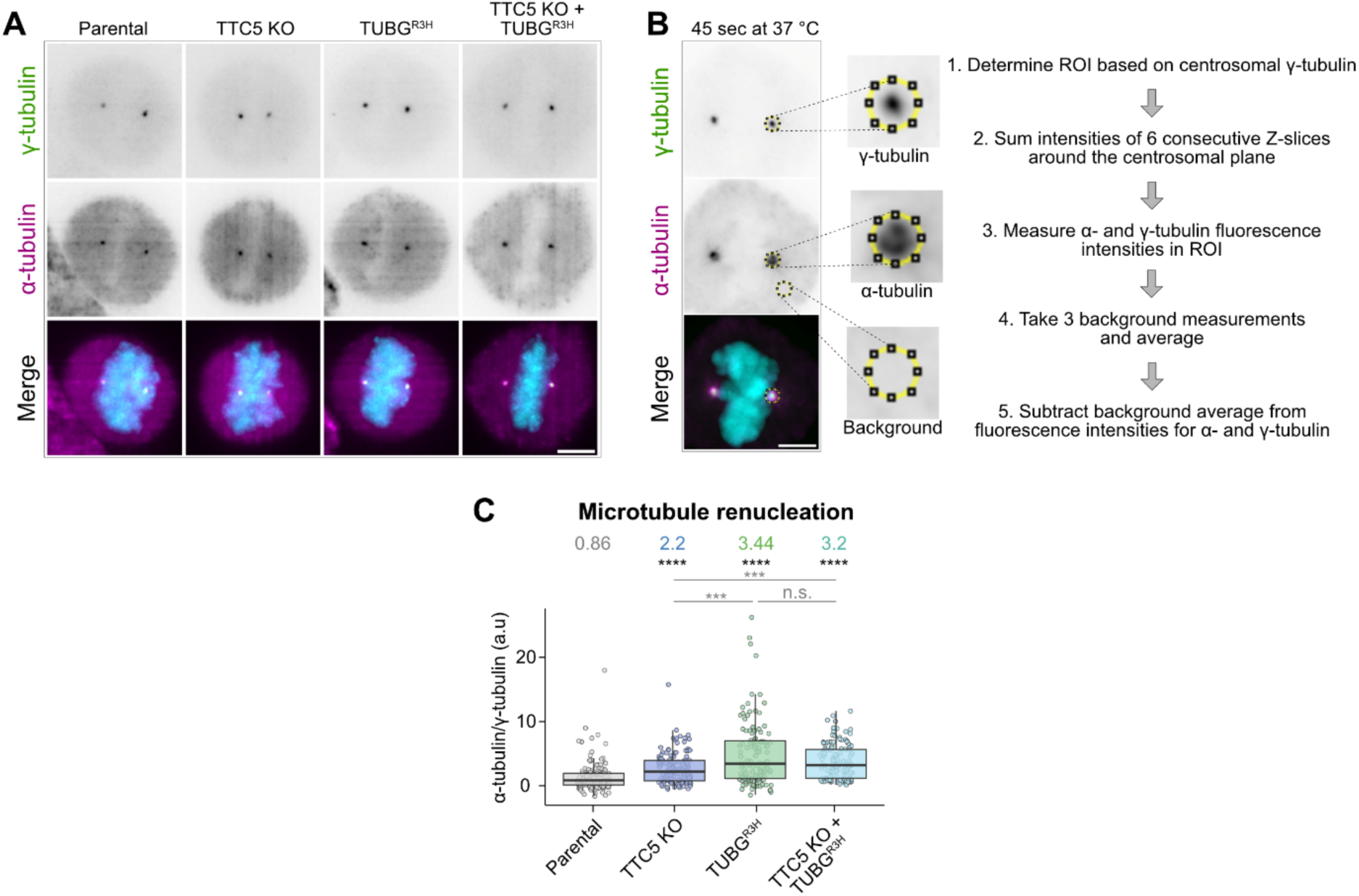
(related to Figure 4). (**A**) Maximum intensity projections of representative images of mitotic HeLa parental, TTC5 KO, TUBG^R3H^, and TTC5 KO + TUBG^R3H^ cells following cold-induced microtubule depolymerisation. Cells were immunostained for γ-tubulin (green) and α-tubulin (magenta). Scale bar = 5 µm. (**B**) Representative image of microtubule regrowth assay, and quantification method performed in Fig. 4A-B. (**C**) Quantification of microtubule re-nucleation analysed in Fig. 4B represented as the α-tubulin/γ-tubulin ratio. Data show the median ± interquartile range. Median values colour-coded by genotype are indicated above the respective boxplots. Shown are Benjamini-Hochberg-adjusted *p-*values from an unpaired Mann-Whitney test comparing ranks for each of the indicated cell lines with the parental cell line as reference, or across the groups. *** *p* < 0.001, **** *p* < 0.0001, n.s. = not significant.

**Figure S7.**
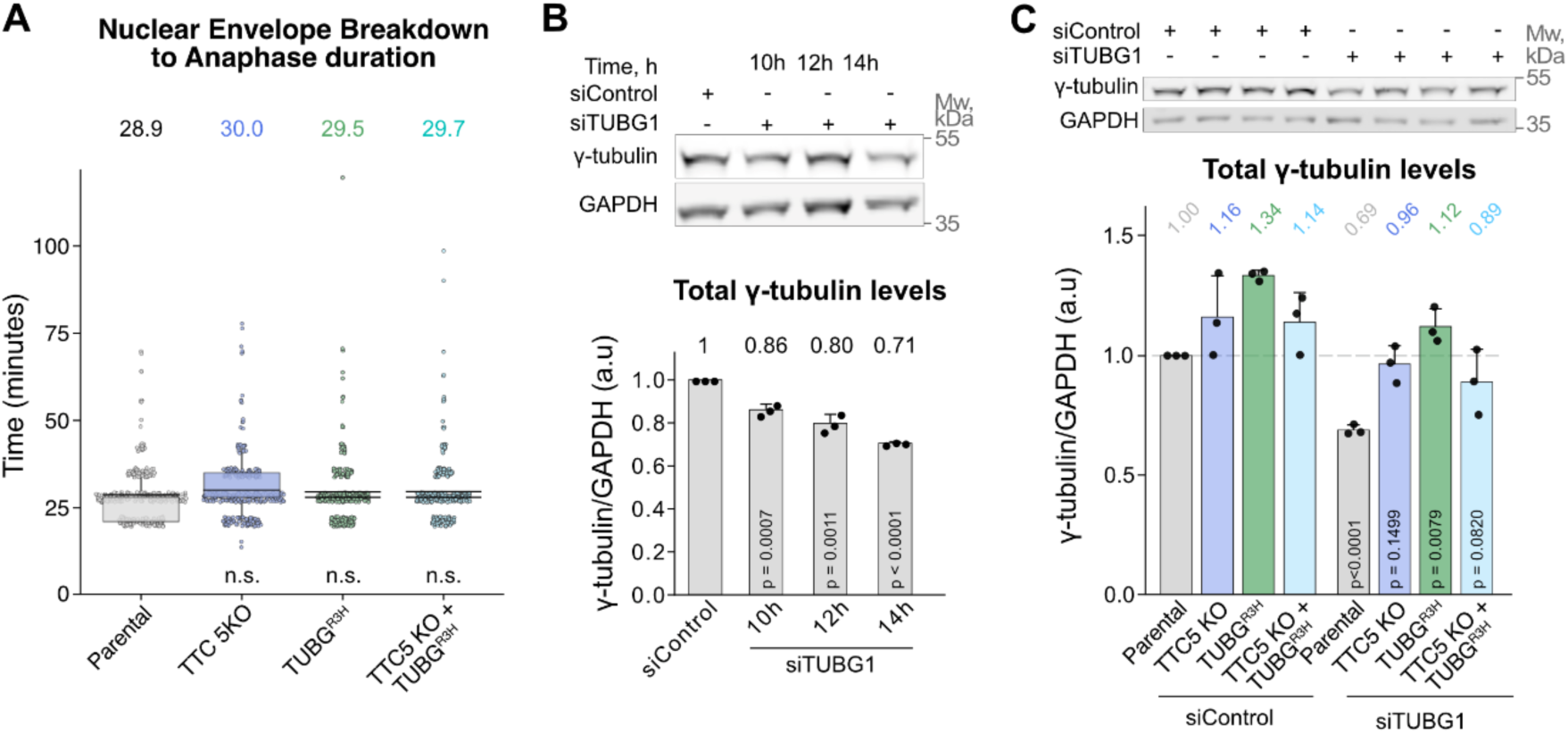
(related to Figure 5). (**A**) Nuclear envelope breakdown to anaphase duration of mitotic cells analysed in Fig. 5A-B. Average mitotic duration is depicted at the top of the panel. Shown are Benjamini-Hochberg-adjusted *p-*values from an unpaired Mann-Whitney test comparing ranks for each of the indicated cell lines with the parental cell line as reference. N.s. = not significant. (**B**) Western blot analysis of total γ-tubulin protein levels in HeLa parental cells treated with siTUBG. Quantification shows γ-tubulin intensity normalized to GAPDH loading control and parental cell line. Data show mean ± SD from three independent experiments, with median values above corresponding bars. Indicated are *p*-values in unpaired, two-tailed Student’s *t*-tests for each treatment with siControl as reference. **(C)** Western blot analysis of total γ-tubulin protein levels across the indicated cell lines treated with siControl or siTUBG, analysed in Fig. 5A-B. Quantification shows γ-tubulin intensity normalized to GAPDH loading control and parental cell line. Data show mean ± SD from three independent experiments. Indicated are *p*-values in unpaired, two-tailed Student’s *t*-tests for each genotype, with the corresponding siControl genotype as a reference.

